# Gut microbiota promote early-life digestive function in songbirds

**DOI:** 10.64898/2026.07.09.736762

**Authors:** Brian K. Trevelline, Jennifer L. Houtz, Catherine R. Andreadis, Jon G. Sanders, Megan K. Collins, Natalie J. Morris, Tosha R. Kelly, Melissah Rowe, Andrew H. Moeller

## Abstract

The vertebrate digestive tract harbors complex microbial communities whose influences on phenotypes and fitness in non-model organisms remain poorly understood. Here, we show that colonization with gut microbiota is critical for digestive development and function in a non-model songbird, the House sparrow (*Passer domesticus*). We raised nestlings under sterile (axenic) conditions and compared them to nestlings reconstituted (conventionalized) with microbiota from adult sparrows. Conventionalization drove significant increases in villus length, crypt depth, goblet cell density, and mucosal thickness in the small intestine. Conventionalized nestlings also exhibited increased growth of several digestive organs and elevated circulating bile acid levels, including bacterial metabolites known to promote growth and lipid metabolism in vertebrates. These results reveal functions of songbird microbiota and establish axenic methods for non-model oviparous vertebrates.

## Main Text

Animals ubiquitously associate with microbial assemblages, collectively termed microbiota, which can affect host phenotypes and fitness through interactions ranging from mutualism to antagonism (*1*). Manipulations in hosts reared under axenic (*i.e.,* sterile) conditions provide the best available approach for quantifying the phenotypic effects of microbiota, enabling mechanistic studies of host-microbiota interactions (*2–7*). In vertebrates, axenic experiments have historically required expensive facilities and equipment available for only a limited number of laboratory and agricultural model organisms. This work has demonstrated host dependence on microbiota for the development of phenotypes critical to host fitness (*1, 8–10*), but the extent to which these findings translate across vertebrate species is unclear (*11–15*). For example, comparative studies of songbirds (Passeriformes), the most ecologically successful and diverse order of terrestrial vertebrates, have shown that these hosts harbor low microbial biomass and highly variable gut microbiota characterized by a relatively weak association with host phylogeny compared to mammals, suggesting that songbirds may have lost reliance on their gut microbiota (*14, 16*). Controlled experiments to test the effects of microbiota in songbirds—and, more generally, in non-model oviparous (egg-laying) vertebrates that represent most terrestrial vertebrate species—have been limited by the lack of robust, low-cost axenic approaches.

Here, we developed an axenic experimental system for oviparous vertebrates (Materials and Methods) and applied it to quantify phenotypic effects of gut microbiota on one of the most abundant and widely distributed wild bird species (*17*): the House sparrow (*Passer domesticus*). House sparrow eggs were collected from a wild-derived captive breeding colony maintained at the Netherlands Institute of Ecology soon after laying and externally sanitized by immersion in a commercially available peracetic acid disinfectant followed by UV-irradiation (*18*). Sanitized eggs were incubated under sterile conditions in an automatic egg incubator and transferred to sterile HEPA-filtered cages prior to hatch. Cages were housed in a custom incubator equipped with automated temperature and humidity control (Materials and Methods; *19*). Nestlings were randomly assigned to either axenically-raised (AX; n = 15) or conventionalized (CV; n = 13) treatment groups upon hatch, with all nestlings receiving a standardized, age-specific volume of sterile formula developed specifically for House sparrows (*20*) delivered by micropipette approximately every 30–60 minutes for 14 hours per day for 7 days (data S1; Materials and Methods). Every other feeding for the first 7 hours during each day from hatch (day 0) to 6 days post-hatch, CV nestlings received formula containing an inoculum derived from homogenized adult House sparrow feces, while AX nestlings received a sham inoculation of sterile deionized water. The meconium (the first fecal sample after hatch) was collected from nestlings shortly after hatch, and subsequent fecal samples were collected daily under direct observation a minimum of 3 hours following the CV inoculations to eliminate the possibility of detecting transient bacteria from the microbial transplants. At 7 days post-hatch, nestlings were fed sterile formula every 30-60 minutes for approximately 2-4 hours prior to dissection to quantify microbial communities of the small intestine contents, blood metabolites, and internal digestive phenotypes. Small intestine contents from age-matched nestlings (7 days post-hatch; n = 10) raised by their parents in captivity were also collected to confirm that reconstitution of gut microbiota approximated gut-microbiota composition of naturally raised nestlings. This experimental design allowed us to test the effects of early-life microbial colonization on developmental phenotypes in sparrow nestlings.

## Validation of axenic rearing

We began by determining whether the axenic rearing of nestlings effectively precluded the establishment of gut microbiota, as well as whether inoculating nestlings with fecal material from adult sparrows led to successful colonization of gut microbiota. To validate these approaches, we conducted amplicon sequencing and qPCR of the 16S rRNA gene to quantify relative bacterial abundances and bacterial load from the intestinal contents and fecal samples of AX and CV nestlings collected over the course of nestling development, the inocula used for microbial transplants, and several negative control samples. These negative controls allowed us to account for possible microbial DNA contamination from commercial extraction kits, sample storage buffer, sterile formula, autoclaved deionized water, and sterile collection swabs (data S2; Materials and Methods). We first compared the composition of AX and CV nestling microbiota in small intestine contents and fecal samples, the inocula used for conventionalization, and several blank extraction controls using Robust Aitchison PCA of 16S rRNA gene amplicon sequences. These analyses were conducted before filtering of possible contaminants present in blank extraction controls to enable comparisons of AX and CV nestling microbiota to bacterial DNA present in commercial extraction kits. This analysis revealed significant differences in microbiota composition in all pairwise comparisons between groups (PERMANOVA Pseudo-F = 19.63, P = 0.001; Fig. 1A; fig. S1; data S2). Gut microbiota of AX nestlings were significantly more similar on average to blank extraction controls than were gut microbiota of CV individuals (t = 6.54; P < 0.0001; fig. S2A). Similarly, microbiota of CV nestlings were significantly more similar on average to microbiota present in the fecal inoculations than were microbiota of AX individuals (t = 6.40; P < 0.0001; fig. S2B). Moreover, gut microbiota of CV individuals were significantly more similar on average to microbiota present in their specific inoculum than to microbiota in other fecal inocula (t = 2.03, P = 0.045; fig. S3). These results confirmed that AX rearing precluded the establishment of gut microbiota in AX nestlings, whereas the CV nestlings harbored microbiota derived from the inocula containing adult sparrow gut microbiota.

**Fig. 1.**
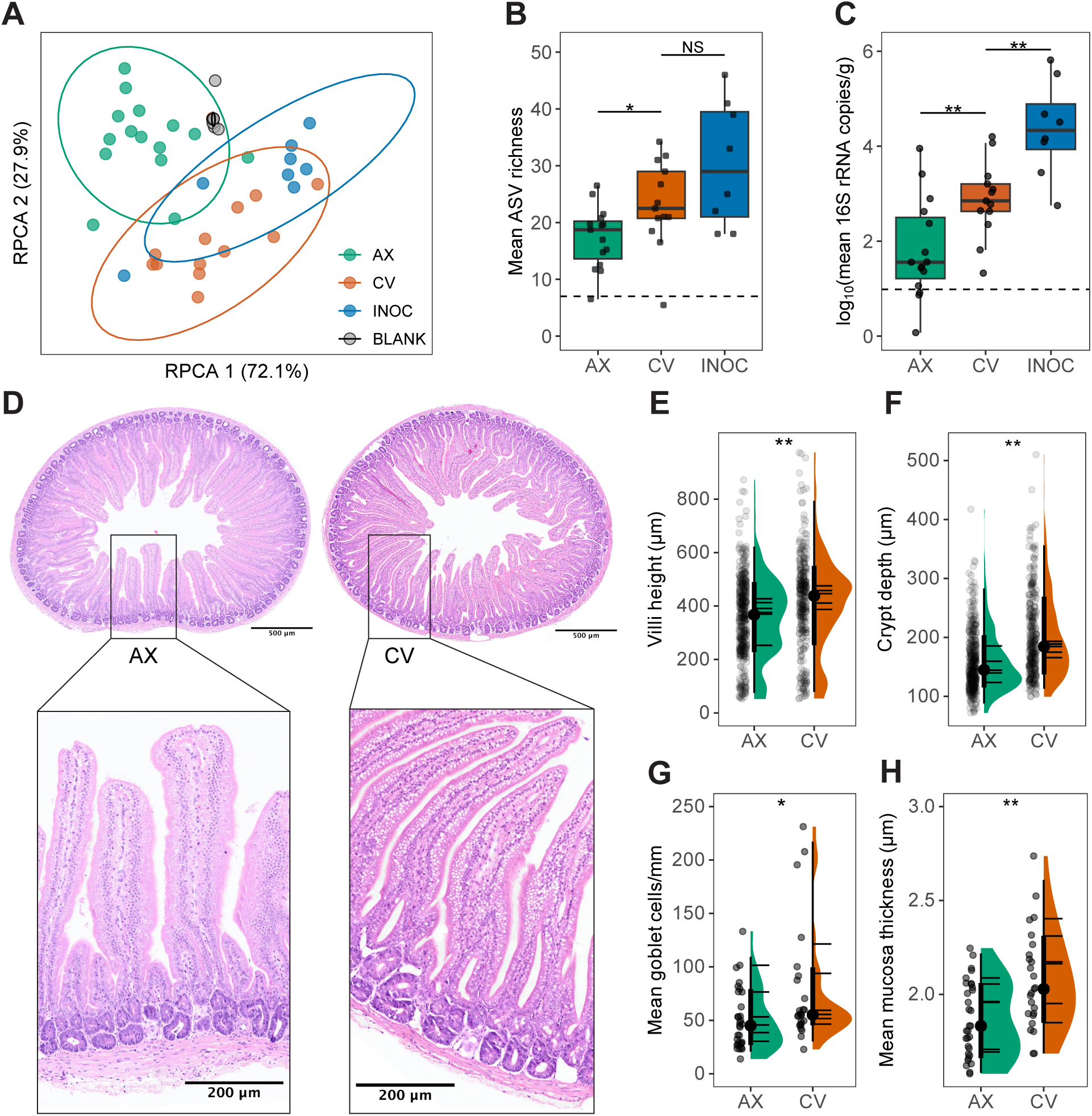
The gut microbiota promotes early-life intestinal development in songbirds. (**A**) Robust Aitchison PCA (RPCA) ordination of bacterial 16S rRNA amplicon sequence variants (ASVs) shows significant differences in microbial community composition between groups with 95% confidence ellipses around group centroids. Axenically-raised (AX; n = 15) showed similar community compositions to blank extraction controls (BLANK; n = 14), whereas conventionalized nestlings (CV; n = 13) showed similar community compositions to inoculated microbial communities (INOC; n = 8). (**B**) Boxplots show that AX nestlings exhibited significantly reduced ASV richness compared to CV individuals, while there was no difference in richness between CV individuals and the inocula used for conventionalization. ASV richness values from AX nestlings overlapped with the maximum observed in negative control samples (dotted horizontal line). (**C**) Boxplots show that the intestinal contents of AX individuals contained significantly fewer copies of the 16S rRNA gene compared to CV individuals.16S rRNA gene amplicon copy numbers observed in AX individuals overlapped with those observed in negative control samples (dotted horizontal line). In (**B**) and (**C**), significance of linear mixed effects models accounting for parentage and sex is indicated by asterisks; P ≤ 0.05 *; P ≤ 0.01 **; P ≥ 0.05 NS. (**D**) Image shows comparison of representative small intestine cross-sections and villi morphology for AX (n = 6) and CV (n = 6) nestlings. Violin plots of morphological measurements from AX (n = 42) and CV (n = 32) nestling small intestine cross-sections at 7 days post-hatch show that CV nestlings had increased **E**) villi height, (**F**) crypt depth, (**G**) goblet cell density, and (**H**) thickness of the mucosa compared to AX nestlings. In (**E–H**), gray points represent individual measurements, black points and bars indicate mean and standard deviation, and horizontal lines within violin plots indicate median values for individual nestlings. Significance of linear mixed effects models accounting for parentage, sex, and individual is indicated by asterisks; P ≤ 0.05 *; P ≤ 0.01 **; ≤ 0.001 ***.

After filtering ASVs detected in the negative controls, we found that the ASVs that successfully colonized CV nestlings were predominantly bacteria in the Order Lactobacillales–Lactobacillus (26.4%), Catellicoccus (21.9%), and Enterococcus (9.3%)–resulting in CV nestling gut microbiota that approximately reflected the genus-level relative taxonomic composition of the microbial inoculations (fig. S4A; data S3). Nearly half (mean of 46.3% ± 21.7% SD) of the ASVs present in fecal inocula were detected in the feces of their specific recipient CV nestling 24–48 hours after receiving their first inoculation (fig. S4B), indicating that these bacteria were successfully transferred to the recipient CV nestlings. To further confirm the efficacy of microbiota conventionalization, we conducted additional 16S rRNA gene sequencing of the intestinal contents of age-matched sparrows raised by parents in captivity. Analyses of these data in the context of experimental sparrows indicated that gut microbiota of CV and parent-raised sparrows exhibited similar genus-level microbiotas compositions (fig. S4A), further confirming successful microbiota conventionalization in nestlings. Together, these results indicate that axenic rearing is an effective approach for precluding the establishment of sparrow gut microbiota of nestlings, and that inoculation with adult fecal material can be used to colonize the gut microbiota for manipulative experiments.

We next tested the effects of AX and CV treatments on gut microbiota richness and absolute abundance. 16S rRNA microbial inventories from fecal samples and intestinal contents demonstrated that AX nestlings had significantly reduced ASV richness compared to CV individuals (t = 2.388, FDR-adjusted P = 0.037, Fig 1B), and this effect remained significant when accounting for nestling sex and parentage (t = 2.321, P = 0.029). In contrast, there was no difference in ASV richness between CV nestlings and the fecal inocula (t = 1.434, FDR-adjusted P = 0.155; Fig. 1B). Intestinal contents of AX individuals contained significantly fewer copies of the 16S rRNA gene compared to CV individuals (t = 2.92, FDR-adjusted P = 0.006, Fig. 1C). This difference remained significant after using linear mixed-effects models that accounted for sex and parentage (t = 6.21, P = 0.008; data S4). Moreover, the distributions of bacterial abundances and ASV richness from AX nestlings overlapped with the maximum abundance observed in blank extraction controls that contained no bacteria (Fig. 1B, C). Interestingly, we observed 207 ASVs in the feces of AX nestlings 24-48 hours post-hatch, and 154 of these ASVs were identified as the genus Lactococcus (mean of 67.2%; fig. S4). This observation suggests that some bacteria may be transmitted in the egg before hatching (*21*). To test for *in ovo* transmission, we profiled 16S rRNA gene diversity in the meconium of AX and CV nestlings collected prior to any feedings or inoculations. Because the meconium forms before hatching, we reasoned that ASVs present in meconium samples would represent bacteria transmitted from mother to offspring through the egg during embryonic development. We detected a total of 91 ASVs in nestling meconium samples (n = 15) that were not detected in any contamination control samples, 11 of which were also identified in AX nestling feces collected 24-48 hours after hatch (n = 14). Among the 11 shared ASVs, 8 were identified as the genus Lactococcus (7 ASVs identified as L. lactis), which dominated the AX nestling microbiota (mean of 67.3%; fig. S4A) at near identical proportions to those observed in meconium samples, indicating that these taxa may have been vertically transmitted *in ovo*. *L. lactis* is commonly used as a probiotic to enhance domestic poultry growth (*22*) and has been previously associated with accelerated development of House sparrow nestlings (*23*). Together, these results indicate that a subset of gut bacterial species in songbirds were transmitted vertically *in ovo*, but that axenic rearing effectively impeded the post-hatching transmission of bacteria and limited the assembly of a gut microbiota in nestlings.

## Microbiota influence gastrointestinal development

We next assessed how conventionalization of sparrows with gut microbiota after hatching affected the development of the gastrointestinal tract, which provides the primary interface for interactions between gut bacteria and the host. Previous work has shown that epithelial differentiation in the small intestine depends on the presence of gut bacteria in germ-free mice (*24*), zebrafish (*9*), and domestic poultry (*25*), but it is not clear whether this dependency also exists in wild songbirds, which may have lost close associations with dense and complex gut microbiota (*14*). To address this knowledge gap, we tested whether CV nestlings exhibited advanced development of the intestinal mucosa in the small intestine, which is responsible for host nutrient absorption (*26*), compared to AX individuals; as would be expected if gut microbiota play critical roles in growth and development of the digestive system in the early life of songbirds. To quantify intestinal development, we conducted detailed morphological measurements for several histological markers of intestinal mucosa development (*25, 27, 28*) from stained tissue sections of the small intestine of 6 individuals from each treatment group (Fig. 1D; fig. S5; Materials and Methods; data S5). We then used mixed-effect linear models with a fixed-effect of sex and random effects for individual (to control for repeated measures) and parentage to test for significant differences in histological markers of intestinal development between CV and AX nestlings. Strikingly, we found that CV sparrows had a significantly more developed small intestine compared to AX sparrows characterized by increased villi height (t = 3.56, P = 0.007; Fig. 1E), deeper intestinal crypts (t = 3.93, P = 0.007; Fig. 1F), increased density of mucus-secreting goblet cells (t = 2.10, P = 0.046; Fig. 1G), and thicker intestinal mucosa (t = 4.56, P = 0.001; Fig. 1H). In contrast, there was no effect of treatment on villi width (t = 1.892, P = 0.092; fig. S6). The effects of the microbial inoculations on the differentiation of intestinal villi suggests that the gut microbiota increased the capacity for host nutrient absorption (*25, 26*), the primary determinant of songbird nestling growth and development (*29*). These results demonstrate that the presence of gut microbiota is essential to the early-life development of intestinal tissue in songbird nestlings.

Given the effects of microbiota conventionalization on the differentiation of intestinal tissues, we next tested whether AX and CV sparrows differed in the development of organs involved in host digestion and nutrient metabolism. Using linear mixed-effects models with tarsus length and sex as covariates, and random effects of parentage, we found that microbiota inoculations into axenically-raised nestlings significantly increased the size of the liver (F = 5.16; P = 0.031; Fig. 2A) and large intestine (F = 4.96, P = 0.035; Fig. 2B), with a trend towards positive effects on small intestine (F = 2.34, P = 0.138; Fig. 2C) and pancreas mass (F = 2.30, P = 0.141, Fig. 2D; data S6). We observed no effect of microbial inoculations on gizzard and heart masses (fig. S7), morphometric measures of avian growth (body mass, tarsus length, and wing chord; fig. S8), or nestling growth and feeding rates (fig. S9). These results indicate that the gut microbiota promotes the development of the digestive system to influence host phenotypes involved in nutrient absorption.

**Fig. 2.**
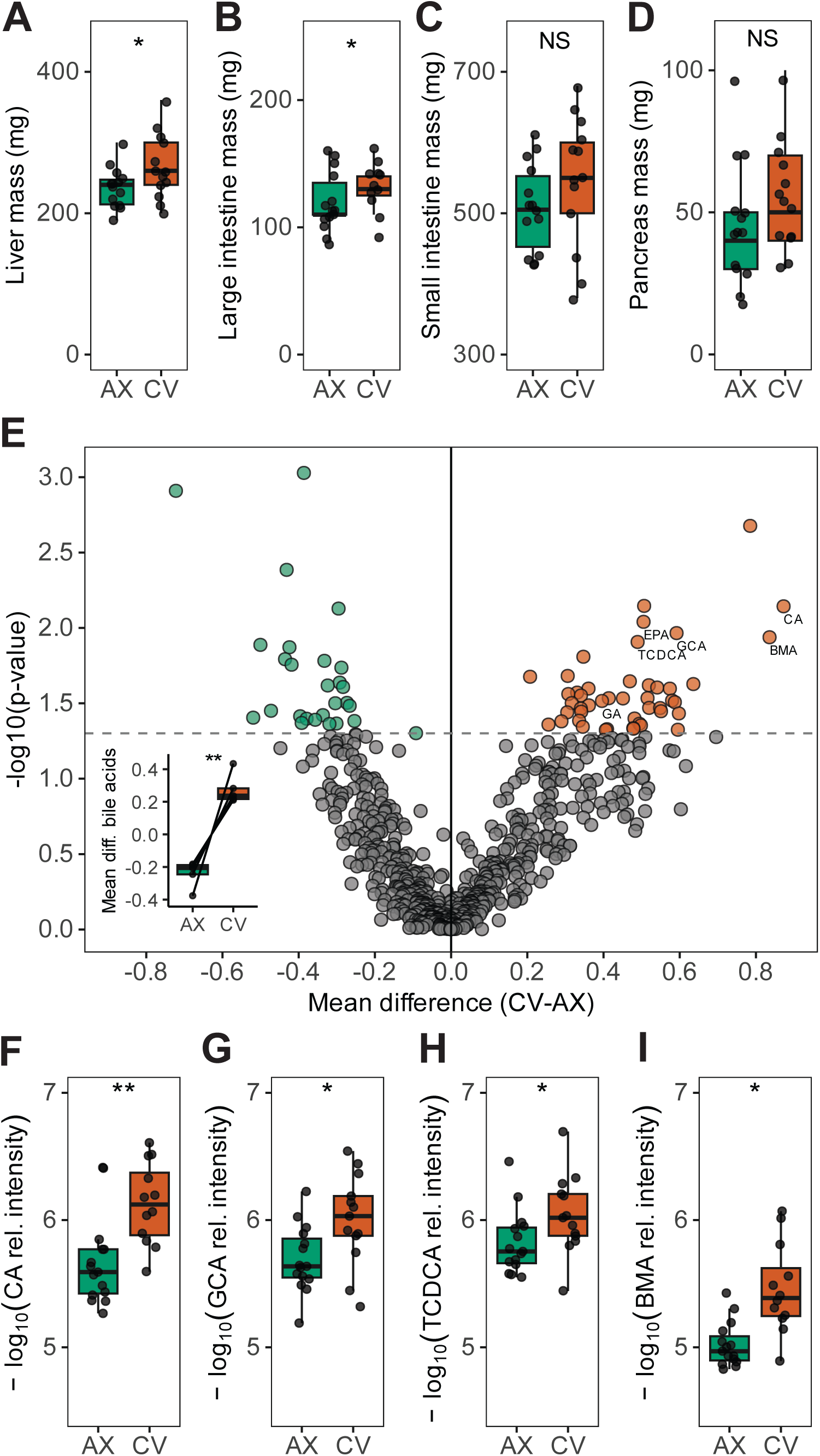
The gut microbiota modulates early-life digestive and metabolic phenotypes in songbird nestlings. Boxplots of digestive organ masses measurements at 7 days post hatch show that microbiota inoculations into axenically-raised nestlings had significant positive effects on the growth of the (**A**) liver and (**B**) large intestine, with a trend towards positive effects on (**C**) small intestine and (**D**) pancreas mass. In (**A–D**), asterisks denote significance of differences between AX and CV organ masses based on linear mixed effects models accounting for tarusus length, parentage, and sex; P ≤ 0.05 *; P ≤ 0.01 **; P > 0.05 NS. (**E**) Volcano plot of plasma metabolites indicate that CV nestlings had significantly higher levels of the primary bile acids: cholic acid (CA), glycocholic acid (GCA), taurochenodeoxycholic acid (TCDCA), and β-muricholic acid (BMA) compared to AX nestlings. Red points indicate metabolites displaying higher relative abundance in CV than in AX nestlings, green points indicate metabolites with higher relative abundance in AX than in CV nestlings, and dashed line indicates significance threshold of P = 0.05 (distributions of p-values shown in fig. S11). (**E, inset**) Boxplot shows all bile acid metabolites (pairs of points connected by lines) detected by RPLC-MS/MS in negative ion mode, and asterisk indicates significance of paired t-test for differences in mean relative peak intensity of bile acid metabolites between CV and AX; P ≤ 0.01 **. Boxplots show significant elevation of log_10_-scaled relative peak intensities of bile acids in CV nestlings: (**F**) cholic acid (CA), (**G**) glycocholic acid (GCA), (**H**) taurochenodeoxycholic acid (TCDCA), and (**I**) β-muricholic acid (BMA). Significance based on linear mixed effects models accounting for parentage is indicated by asterisks; FDR-adjusted P ≤ 0.05 *, P ≤ 0.01 **.

## Microbiota influence nestling metabolomes

To further interrogate how gut microbiota influenced the processing of nutrients in the early life of songbirds, we next tested whether microbial inoculations altered levels of circulating plasma metabolites in nestlings. We used RPLC-MS/MS (for hydrophobic lipids and fatty acids) and HILIC-MS/MS (for hydrophilic peptides and sugars) to quantify the relative peak intensities of circulating plasma metabolites in AX and CV nestlings. For each chromatography and polarity type, we used FDR-corrected linear mixed effects models that accounted for host parentage to test for metabolites that differed significantly in relative abundance between AX and CV birds (data S7). Metabolomic profiles of AX and CV birds were distinguishable in multivariate compositional space across all chromatography and polarity types (fig. S10), and distributions of p-values from linear mixed effects models displayed a greater number of significant (P ≤ 0.05) differences beyond that expected under the null hypothesis (*i.e.,* > 5% of tests), as indicated by the rightward skews of the distributions of p-values for each mode (fig. S11; data S7). These results demonstrate that microbial inoculations had broad effects on nestling metabolism. Analyses of the annotation categories to which differentially abundant metabolites belonged indicated that differences between metabolite profiles of AX and CV nestlings were primarily driven by compounds involved in host digestion and lipid metabolism (data S7; Fig. 2E). We next tested whether the metabolites annotated within specific categories displayed consistent differences between AX and CV nestlings. For these analyses, we used paired t-tests to determine whether the mean metabolite relative abundance within each annotation category differed significantly between CV and AX nestlings, finding that only the ‘bile acids’ category differed significantly between treatment groups at the FDR-adjusted P ≤ 0.05 threshold (Fig. 2E; data S7). Specifically, each of the metabolites annotated as bile acids displayed elevated relative abundances in CV compared to AX nestlings, and annotations belonging to the ‘bile acid’ category displayed significantly higher mean abundances in CV compared to AX nestlings (t = 6.671, FDR-adjusted P = 0.0026; Fig. 2E, inset; data S7). Results of linear mixed effects models of individual metabolites indicated that CV nestlings exhibited significantly higher levels of the primary bile acids cholic acid (t = 3.053, FDR-adjusted P = 0.007; Fig. 2F), glycocholic acid (t = 2.861, FDR-adjusted P = 0.011; Fig. 2G), taurochenodeoxycholic acid (t = 2.797, FDR-adjusted P = 0.012; Fig. 2H), and β-muricholic acid (t = 2.831, FDR-adjusted P = 0.012; Fig. 2I) compared to AX individuals. In vertebrates, primary bile acids are synthesized and conjugated in liver hepatocytes before secretion into the small intestine, where they are essential for the absorption of dietary lipids (*30*). In the small intestine, primary bile acids are modified by microbiota into secondary bile acids, which promote cellular proliferation of intestinal villi (*31, 32*) and bacterial bile acid modifications in the gut have been shown to regulate host growth and lipid metabolism in mice (*33*). In addition to bile acids, CV nestlings exhibited significantly higher levels of other metabolites known to promote intestinal cellular proliferation and barrier function, including eicosapentaenoic acid (*34*) and glyceric acid (*35*) (fig. S12). Overall, these results indicate that the presence of the gut microbiota in early life promotes diverse blood-circulating metabolites with known roles in intestinal development and metabolism, providing a possible mechanistic basis for the effects of microbiota on host intestinal cellular proliferation (Fig. 1D–H) and the growth of digestive organs (Fig. 2A–D).

## Conclusions

In this study, we demonstrated that the gut microbiota promotes the early-life development and function of the digestive system in songbirds, spanning levels of biological organization including cellular development in the intestine, the growth of digestive organs, and the blood metabolome. Contributions of the gut microbiota on digestive development may be mediated by microbiota effects on lipid and bile acid metabolism, as evidenced by the significant effect of microbial colonization on host blood metabolite profiles. More broadly, the axenic methods that we developed can be used to conduct controlled microbiota depletion and colonization experiments on any oviparous terrestrial vertebrate whose hatchlings can be reared in captivity. These approaches set the stage for experimental studies of microbiota-host interactions in a diversity of non-model host species.

## Supporting information

dataS1

dataS2

dataS3

dataS4

dataS5

dataS6

dataS7

## Acknowledgments

We thank NIOO staff Marcel Visser, Christa Mateman, Judith Risse, Kees van Oers, Agata Pijl, Anne Dijkzeul, Veronika Hohnec, Djan Mattijssen, Philipp Ratasiewicz, Kristen Rosamond, Lindy Schneider, Tessie van der Honing, and Ronald van Beek. We thank Cornell personnel Irby Lovette, Maren Vitousek, Andy Lui, Weiwei Yan, Mary-Margaret Ferraro, Bronwyn Butcher, Riley Antony, Alice Hu, Sarah Tang, and Julia McCabe. We thank Bill Karasov and Enrique Cadieves-Vidal for their guidance in the development of the nestling feeding protocols. We thank the Brain Health Research Institute Neuroimaging Collaboratory at Kent State University for access to resources used in the analysis of histology data.

## Author contributions

Conceptualization: BKT

Formal Analysis: BKT

Methodology: BKT, JLH, CRA, JGS, MKC, MR, AHM

Investigation: BKT, JLH, CRA, MKC, MJM, TRK, MR, AHM

Visualization: BKT

Funding acquisition: BKT, MR, AHM

Project administration: BKT, MR, AHM

Supervision: BKT, MR, AHM

Writing – original draft: BKT

Writing – review & editing: BKT, JLH, CRA, JGS, MKC, MJM, TRK, MR, AHM

## Data, code, and materials availability

All data, code, images and incubator details are publicly available in Zenodo (*36*). Raw sequence data is publicly available in the NCBI Short Read Archive (SRA) under BioProject accession PRJNA1477154l.

## Materials and Methods

### Axenic rearing

Fertilized eggs were sourced from a captive population House sparrows (*Passer domesticus*) at the Netherlands Institute of Ecology (NIOO-KNAW, Wageningen, NL). Each aviary housed 6-8 wild-caught adult sparrows (at equal sex ratios) and nestboxes to support breeding for up to 4 nesting pairs. Eggs were collected approximately 1-2 hours after laying and were removed from nests by hand with ethanol-sanitized gloves and immediately transferred to a sterile laminar-flow cabinet for external sanitization. Eggs were externally sterilized by immersion into 0.075% peracetic acid (1.5% Divosan Activ®; Diversey) at 40°C for 15 seconds, rinsed briefly in sterile deionized water at 40°C, followed by UV-C (254 nm) irradiation using an ultraviolet sterilizer (Phillips) at approximately 900 μJ/cm2 for 10 seconds (*18*). Sanitized eggs were then transferred to an automatic egg incubator (Brinsea Maxi 24 EX) sanitized using 0.075% peracetic acid, 70% ethanol, and UV-irradiation and housed in a dedicated sterile laminar-flow cabinet. Eggs were incubated at 37.5°C and 55% humidity (using sterile deionized water) with a 180° rotation 8-times per day. After 10 days of incubation, eggs were then transferred to a sterile glass dish housed inside a sterile HEPA-filtered cage designed for maintaining germ-free mice (MSX4; Innovive). Each sterile cage was then placed in a custom incubator with an automated forced-air heating and humidification system fitted with a 0.22 µm in-line filter to maintain sterility of the incubation environment (full description and protocols available in Zenodo (*35*). Incubators were maintained at approximately 37.5°C and 55% humidity with an increase to 70% humidity for the final 48 hours of egg incubation.

Hatchlings were weighed within 1 hour of hatch during which the meconium (the first fecal sample after hatch; n = 16) was collected from both AX (n = 7) and CV (n = 8) treatment groups (one sample failed due to 0 reads data after filtering) soon (1-3 hours) after hatch. Meconium samples were collected either from a sterile foil sheet used for weighing or directly from the hatchling using a sterile swab and immediately stored in DNA/RNA Shield (Zymo Research) and then flash frozen on dry ice, before being stored at -80°C for later processing. Hatchlings were transferred to a sterile stainless-steel dish lined with autoclaved cotton wool and tissue paper housed inside a sterile HEPA-filtered cage (MSX4; Innovive) maintained at approximately 37.5°C and 55% humidity. Nestlings were housed either singly or in pairs of unrelated individuals (eggs sourced from different clutches) and fed an age-specific volume of sterile semi-synthetic formula developed specifically for House sparrows (*19*) beginning at the onset of begging behavior (approximately 1-3 hours post-hatch). The gamma-irradiated powdered diet (data S1) was mixed 1:4 with sterile deionized water and stirred for 30 minutes at 45°C. Pre-warmed (40°C) sterile formula was delivered to nestlings by sterile aerosol barrier filter wide-bore micropipette tips (P200 and P1000 from Corning and Rainin) approximately every 30 minutes for 14 hours per day for 7 days. Feeding volumes were approximately the following age-specific volumes: Day 0: 20-60 µL (increasing in 10 µL increments every 4-6 hours depending on nestling hatch mass and condition); D1: 50-60 µL (based on nestling condition); D2: 70–80 µL; D3: 95–135 µL; D4: 140 µL; D5: 170 µL; D6: 210–215 µL; D7: 240 µL. For these feedings we aimed to standardize volumes to ensure equal levels of food intake across treatment groups; however, small adjustments in feeding volume (within 10%) were made based on feeding motivation of the nestlings. Post-hoc tests confirmed that there was no significant difference in AX and CV nestling food intake (fig. S9B). Nestling body mass was measured three times daily during the first, middle, and last feeding of the day that were averaged to calculate mean daily body mass. All experimental work was conducted by wearing full PPE (gowns, hair nets, shoe covers, surgical masks, and gloves sanitized with 70% ethanol) in a dedicated clean room equipped with a HEPA-filtered air handler and was regularly sanitized using Virkon S disinfectant (Lanxess). All fecal sample collections, morphological measurements, and feedings were conducted under sterile conditions in a laminar-flow cabinet. All animal experiments were performed at the Netherlands Institute of Ecology (NIOO-KNAW) under license from the Central Authority for Scientific Procedures on Animals (CCD Project AVD-80100 2022 15876 to MR) and approved by the Animal Welfare Body (IvD) of the Royal Netherlands Academy of Sciences (KNAW; protocol number NIOO22.11).

### Microbial transplants

Nestlings were randomly assigned to either axenically-raised (AX; n = 19) or conventionalized (CV; n = 14) treatment groups upon hatch, with clutch-mates split across treatment groups to minimize potential paternal effects on nestling phenotypes. Inocula for nestling conventionalization were derived from sterile-collected fecal samples from 6-8 captive adult House sparrows (at equal sex ratios) occupying unique aviaries (n = 8) as inocula to mimic natural microbiome variation in nature. Fecal sampling for microbial inoculum was performed using a sterile sample collection method, modified slightly from Knutie and Gotanda (2018). Specifically, each bird was placed into a clean flat-bottomed wooden box containing a sterile collection tray fitted with a wire mesh platform, this allowed fecal material to fall to the sterile surface and prevented any contact with the fecal material. All collection materials were sterilized prior to use, first by submersion in a 10% bleach bath the night before use and then with 70% ethanol spray immediately before sampling. Following defecation, fecal samples were transferred from the collection tray to a sterile microcentrifuge tube using a sterile spatula. Fecal material from adult sparrows in each aviary were pooled and homogenized in sterile water to achieve a concentration of 0.75 mg/µL (dry weight/vol) that were aliquoted into 10 µL doses and frozen at -80°C for less than 2 weeks prior to inoculation into conventionalized nestlings. Frozen inocula aliquots were resuspended in sterile formula and delivered to conventionalized nestlings once per hour for the first 7 hours of feedings on days 0-6 post-hatch, while AX nestlings received a sham inoculation of 10 µL sterile water. Daily fecal samples were collected from a sterile foil sheet on which nestlings were placed to defecate using a sterile swab during nestling measurements and immediately stored in DNA/RNA Shield (Zymo Research), flash frozen on dry ice, and stored at -80°C for approximately 6 months prior to DNA extraction. Fecal samples used for microbial analysis collected a minimum of 3 hours following the CV inoculations to eliminate the possibility of detecting transient bacteria from the microbial transplants.

AX (n = 15) and CV (n = 13) nestlings that successfully reached 7 days post-hatch were sedated using isoflurane inhalation, after which nestlings were weighed using a digital balance (Mettler Toledo; readability ± 0.01 g), and measured for tarsus length using digital calipers (Fisher Scientific; readability ± 0.01 mm) and unflattened wing chord length using a wing ruler (Avinet). Nestlings were then euthanized via rapid decapitation under isoflurane sedation, and we collected blood from the trunk of the animal for the analysis of blood metabolites using untargeted metabolomics. Nestlings were then dissected to quantify internal organ masses (wet weight, blotted dry) using a digital balance (Mettler Toledo; readability ± 0.01 g) and for the collection of small intestine contents for 16S rRNA microbial inventories. All dissections were conducted with sterile instruments. Small intestine contents from the pylorus to the colic ceca were collected and immediately stored in DNA/RNA Shield (Zymo Research), flash frozen on dry ice, and stored at -80°C for approximately 6 months prior to DNA extraction. The small intestine was then divided into three portions of roughly similar length (proximal, medial, and distal sections). An approximately 1 cm section of intestinal tissue was collected from the distal end of the proximal portion of the small intestine and fixed in 4% PFA for 24 hours before being transferred to 70% ethanol for storage and later histological analysis. Small intestine contents from age-matched nestlings (7 days post-hatch; n = 10) raised by their parents in captivity were also collected to confirm that reconstitution of the gut microbiota approximated the composition of naturally raised nestlings. All organ mass data (with the exception of liver mass from one AX individual that was identified as statistical outlier using Grubb’s test and thus removed) were analyzed using linear mixed-effects models that included fixed effects for tarsus length measurements taken from all AX and CV nestlings 7 days post-hatch (to account for variation in structural body size) and sex, and a random effect of parentage in R version 4.3.0 (*36*) using lme4 (*37*). All data and code for analyses are available in Zenodo (*35*).

### 16S rRNA microbial inventories

DNA was extracted from bead-homogenized frozen fecal material (3 cycles at 6.3 m/s for 45 seconds each with 4-minute incubation periods between cycles using the Omni International BeadRuptor Elite) using the PowerFecal Pro Kit (Qiagen). Several modifications were made to the standard extraction protocol to improve DNA extraction yields from bird feces: (1) the combination of 200 µL CD2 and 800 µL of CD3 lysis steps, (2) elution of DNA using 50 µL of pre-warmed C6 buffer and incubation at 60°C for 5 minutes, and (3) and a second 50 µL elution using the buffer from the from the first elution to further concentrate the DNA. DNA extractions from intestinal contents, nestling fecal samples, inocula used for microbial transplants, several blank extraction control samples to account for possible microbial contamination during the extraction procedure and microbial DNA present in commercial extraction kits (*38*), sample storage buffer, sterile formula, autoclaved deionized water, and sterile collection swabs (data S2) were conducted under sterile conditions in a class-II laminar-flow biosafety cabinet to eliminate the possibility of microbial contamination. Extracted DNA was amplified and sequenced by the Weill Cornell Medicine Microbiome Core Facility (New York, NY). PCR was used to amplify a portion the V4-V5 region of 16S microbial small subunit ribosomal RNA gene following the Earth Microbiome Project Protocol using primers 515F-Y (GTGYCAGCMGCCGCGGTAA; *39*) and 926R (CCGYCAATTYMTTTRAGTTT; *40*). Amplicon libraries were sequenced using a 250 bp paired-end (v3 reagents) run on an Illumina MiSeq at a target depth of 100,000 reads per sample. A total of 5,312,786 sequencing reads (median of 30,319, mean of 29,680 per sample [n = 179] ± 22,111 SD) were quality filtered, merged, and clustered into 1,730 Amplicon Sequence Variants (ASVs) with DADA2 (*41*) in QIIME2 version 2023.5 (*42*) using default settings. Samples from AX (mean of 38,035 [n = 59] ± 21,440 SD) and CV (mean of 35,687 [n = 52] ± 15,374 SD) nestlings returned a similar number of reads. Taxonomy was assigned to ASVs using the SILVA reference database release 138 (*43*) and a classifier trained to the 515F-Y/926R primer set in QIIME2 using default settings. Clustered ASVs were filtered to exclude non-bacterial sequences (archaea, chloroplasts, eukaryotes, and mitochondria). Contaminant ASVs detected in control samples of Zymo DNA/RNA Shield (n = 5), autoclaved water (n = 4), nestling formula (n = 8), and a sterile swab (n =1) were filtered from the analysis (feature-table filter-features) to control for potential microbial DNA contaminants, reducing our total number of reads to 1,661,949 (mean of 10,453 per sample [n = 159] ± 11,666 SD) and 1,595 ASVs. Unrarefied ASV tables were averaged (rounding up to nearest whole number of reads; feature-table group --p-mode “mean-ceiling”) across three fecal samples for each individual collected on days 1–2, 3, or 5–6 post-hatch, and small intestine contents collected during nestling dissections on day 7 post-hatch for comparisons of alpha diversity using ASV richness and beta diversity using Robust Principal Component Analysis (RPCA) on Aitchison distances (*44*) in QIIME2. For RPCA (Fig. 1A) and ASV richness (Fig. 1B) analyses only, all ASVs detected in meconium samples (i.e., ASVs for which read count in the meconium was > 0) were filtered to eliminate potential effects of in ovo transmission on the apparent efficacy of axenic-rearing and conventionalization. All ASVs detected in blank DNA extraction controls (n = 16) were filtered from subsequent taxonomic and community composition analyses. Linear mixed-effect models, PERMANOVA, and PERMDISP with False Discovery Rate correction were conducted in the R package lme4 and QIIME2, respectively. Raw sequence data is publicly available in the NCBI Short Read Archive (SRA) under BioProject accession PRJNA1477154. Data and code for analyses are publicly available in Zenodo (*35*)

### PCR assays

We measured 16S rRNA gene copy number in genomic DNA from blank extractions, fecal inocula, and small intestine contents from AX and CV nestlings to estimate absolute bacterial abundance. These analyses used the Femto Bacterial DNA Quantification Kit (Zymo Research) for quantitative PCR according to the manufacturer’s instructions. PCR reactions were carried out in triplicate on a QuantStudio 7 Pro Quantitative PCR using Design & Analysis Software v2.7.0 (ThermoFisher Scientific). Mean quantification cycle (Cq) across triplicates (median coefficient of variation 0.92%, range 0.05 - 3.08%) for each sample were used to calculate the quantity of amplified 16S rRNA product using standard curves and estimate 16S rRNA gene copy number according to the manufacturer’s instructions (data S4). Mean 16S rRNA gene copy number in each sample was multiplied by a correction factor calculated as the ratio of amplicon sequencing reads after the removal of contaminant ASVs (controls, formula, DNA extraction blanks) divided by total sequencing reads (after filtering) to account for the effect of microbial contamination on 16S copy number (data S4). The corrected copy number value for each sample was divided by the mass of material used in DNA extractions to calculate mean copy number per gram and transformed on a log10 scale prior to statistical analysis using mixed-effect linear models in the R package lme4 with a fixed effect of sex and a random effect for parentage.

Nestling sex was determined using DNA extracted from red blood cells or liver tissue collected during nestling dissections with the Qiagen DNeasy Blood and Tissue Kit (Qiagen) according to the manufacturer’s instructions. The CHD1 gene, which differs in length between avian sex chromosomes, was PCR amplified using the P2 and P8 primer set using the following protocol (*45, 46*): 3.750 µL nuclease-free water, 4.688 µL 2X KAPA HiFi HotStart ReadyMix, 0.281 µL of forward and reverse primers, and 1 µL of extracted DNA. PCR conditions were as follows: 2 minutes initial denaturation at 94℃, followed by 39 cycles of 30-second denaturation at 94℃, 45-second annealing at 50℃, 45-second extension at 72℃, followed by a final extension phase of 1 minute at 48℃ and 5 minutes at 72℃. PCR products were run on a 2% agarose gel for 120 minutes at 100 V and then imaged to determine nestling sex. Nestlings were identified as males if a single amplified band was present (ZZ genotype) and identified as females if two amplified bands were present (ZW).

### Parentage analysis

DNA was extracted from either toe clippings (collected post-mortem) or feathers collected from nestlings and adult birds from the NIOO captive population using the FavorPrep Tissue Genomic DNA Extraction 96-well Kit (Favorgen) according to the manufacturer’s instructions. We then genotyped all individuals at five highly polymorphic microsatellite loci developed from house sparrows (Pdo1, Pdo3, Pdo6, Pdo10; *47–49*) or other passerine species (Ase18; *50*). PCR products were separated on an ABI3730 automated sequencer and fragment size was assigned with GeneMapper 6.0 (ThermoFisher Scientific). We assigned maternity and paternity using the program CERVUS 3.0 (*51*), which determines which male and female has the highest likelihood of producing a given offspring. Parentage (*i.e.,* genetic family) was determined by overlapping loci with potential parents. Candidate mothers and fathers were restricted to individuals present in the aviary from which the eggs were collected.

### Histology

Preserved small intestine tissue (see dissection details above) from 6 representative nestlings from AX and CV treatment groups (selected randomly after balancing for sex and parentage) were embedded in paraffin and 2-3 µm thick cross-sections were cut using a rotary microtome and stained with haematoxylin-eosin (H&E) by the Veterinary Pathological Diagnostic Centre at Utrecht University (Utrecht, NL). Stained cross-sections were non-sequential (i.e., distance between stained sections was approximately 20µm) to ensure independent cell populations were represented. Mounted tissues were digitized using an Olympus VS200 slide scanner at 40X magnification at the Brain Health Research Institute Neuroimaging Collaboratory at Kent State University (Kent, OH). Raw TIFF files were white-balanced and processed to enhance image quality using Fiji ImageJ version 1.54p (*52*) and all image files are publicly available in Zenodo (*35*). For each intestinal cross-section measured in AX (n = 42) and CV (n = 32) nestlings, each villus with an identifiable base and tip were numbered clockwise and a random number generator was used to select 10 representative villi (AX: n = 418; CV: n = 319) for which to quantify villus height, width, and crypt depth using QuPath version 0.5.1 (*53*). Villus height was measured using the polyline tool, following the lamina propria within the villi (fig. S5). Villus width was calculated by taking the average of five width measurements, at the base, the tip and each quarter of the length, using the line tool in QuPath. Crypt depth was measured from the base of the villus to the beginning of the muscularis mucosae using the line tool (fig. S5). Differences between AX and CV nestlings villi measurements (data S5) were tested using linear mixed-effect models with a fixed effect of sex and random effects for individual (to control for repeated measures) and parentage.

Digitized images of intestinal cross-sections were loaded into a machine learning model (HistoAnalyzer; https://github.com/TrevellineLab/HistoAnalyzer) to quantify the number of goblet cells and mucosal thickness of the villi (excluding samples with less than 5 mm perimeter). These measurements were conducted using the following settings: --mpp 0.25 --band_inner_um 0 --band_outer_um 200000 --lowH_percentile 35 --max_nuc_density 0.15 --vacuole_band_um 150000 --min_obj_um2 20 --max_obj_um2 400 --lumen_pick_mode largest. This method builds a tissue mask for each image, selects the primary lumen (largest or most central), and defines a mucosal band at a user-specified distance from the lumen using microns-per-pixel scaling. The method then uses the hematoxylin channel, Otsu thresholding, and a local density map to estimate nuclei. Within the mucosal band, “mucin” is quantified as pixels with low hematoxylin signal and low nuclear density. Low hematoxylin and low nuclear density were defined by the --lowH_percentile and --max_nuc_density parameters, respectively (--lowH_percentile: how pale or weakly purple-stained a region must be to count as low hematoxylin, based on the pixel’s percentile; --max_nuc_density: the maximum allowed amount of nearby nuclei/cells for a region to count as sparsely cellular, based on the nuclei-positive fraction of pixels within the local window). The method also computes the mucin area fraction, the lumen perimeter (to mm), counts goblet-like vacuoles in a narrow epithelial band with size filters in µm², and skeletonizes mucin touching the lumen to estimate local mucosal thickness statistics (mean/median/90th/max in µm). Goblet-like vacuoles were defined algorithmically as small, low-hematoxylin, low-nuclear-density objects located within a narrow band outside the lumen.

The HistoAnalyzer python script (https://github.com/TrevellineLab/HistoAnalyzer) was implemented by first quantifying mucin-like material from intestinal cross-section images using a custom image analysis pipeline implemented in Python (scikit-image, SciPy). All analyses were performed on the hematoxylin channel extracted from RGB images. All specific parameter choices were selected after testing the workflow with a range of parameters and visual inspection of output images for accuracy. To standardize staining intensity across images, hematoxylin values were rescaled using percentile-based contrast normalization. Specifically, pixel intensities were linearly rescaled such that the 1st and 99th percentiles of the hematoxylin signal mapped to the minimum and maximum output range, respectively. This reduced the influence of extreme outliers and enabled consistent thresholding across samples. A binary mask of nuclei was generated using global Otsu thresholding of the normalized hematoxylin image. Pixels exceeding the Otsu threshold were classified as hematoxylin-rich (putative nuclear material). Small objects (<16 pixels, corresponding to ∼ 8 µm, or approximately 50% the width of a typical goblet cell) were removed to reduce noise (*54*). Local nuclear density was then estimated for each pixel as the fraction of nuclear pixels within a circular neighborhood of radius 20 µm (converted to pixels based on image resolution). This yielded a continuous map of tissue cellularity, where high values correspond to densely nucleated regions and low values correspond to sparsely cellular or acellular regions. All mucin quantification was restricted to a predefined mucosal band adjacent to the intestinal lumen. This band was defined as a region extending from the lumen boundary outward by a specified distance (120 µm). This spatial restriction ensured that measurements were confined to the biologically relevant region in which luminal mucus is expected (*55*). Within the mucosal band, mucin-like regions were identified based on two criteria. First, pixels were classified as low intensity if their value fell below the 30th percentile of hematoxylin intensities within the mucosal band, allowing adaptive thresholding across samples. Second, pixels were required to have local nuclear density below a fixed threshold of 0.15, excluding cellular regions. The final mucin mask was defined as the intersection of the mucosal band with pixels meeting both criteria. The mask was further refined by removing small, connected components (<50 pixels) and applying a binary opening operation (disk radius = 1 pixel) to reduce noise and smooth boundaries. The fraction of the mucosal band occupied by mucin was calculated as:

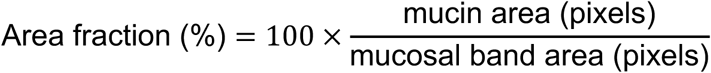

This metric provides a normalized measure of mucus abundance independent of section size. The intestinal lumen was segmented upstream, and its perimeter was computed using region-based measurements. Perimeter length in pixels was converted to millimeters using the known image scale (microns per pixel). This value was used to normalize spatial counts to the length of the mucosal interface. To approximate goblet cell abundance, low-intensity, low-density regions were identified within a narrow band adjacent to the lumen (width 40 µm). Candidate vacuolar structures were defined using the same intensity and density thresholds as above and filtered to remove small objects (<10 pixels). Connected components were labeled, and objects were retained only if their area fell within a specified range (20–400 µm², converted to pixels based on image resolution). The number of such objects was recorded per image and normalized by lumen perimeter (counts per mm). These structures were operationally defined as “goblet-like vacuoles”. To quantify the thickness of the luminal mucus layer, analysis was restricted to mucin regions directly contacting the lumen. Connected components of the mucin mask were identified, and only those overlapping a one-pixel dilation of the lumen boundary were retained. For these regions, a Euclidean distance transform was computed, yielding the distance from each pixel to the nearest boundary. The medial axis (skeleton) of the mucin layer was then extracted. At each skeleton pixel, local thickness was estimated as twice the distance transform value (i.e., diameter) and converted to microns using the image scale. This produced a distribution of local thickness values, from which summary statistics were computed, including mean, median, 90th percentile, and maximum thickness. For each image, an overlay visualization was generated showing the original RGB image with superimposed masks for the mucosal band, lumen, and segmented mucin regions. These overlays were used to visually verify segmentation accuracy. All measurements were computed per image and exported as a table, including descriptive statistics regarding mucosal band area, mucin area, area fraction, lumen perimeter, goblet-like vacuole counts (total and per mm), and mucosal thickness. All statistical analyses of histological measurements were performed using linear mixed-effect models in the R package lme4 (*37*) with sex as a fixed effect and random effects for parentage and individual to account for repeated measures.

### Metabolomics

Blood samples collected from nestlings during dissections were centrifuged at 14,000 rpm for 2 minutes to separate plasma and red blood cells, and frozen at -80°C for later analysis. To perform metabolomics, plasma samples were thawed and proteins were precipitated from 15µL of plasma with 45µL of ice-cold 100% methanol, which was then vortexed for 10 seconds, incubated at 4°C for 1 hour, and centrifuged for at 18,000 xg at 4°C for 10 minutes. The supernatant was transferred to a sterile centrifuge tube, dried-down using a vacuum centrifuge, and shipped frozen to the Proteomics and Metabolomics Core facility at Cornell University (Ithaca, NY) for LC-MS/MS untargeted metabolomic analyses. Precipitates were resuspended in 200µL of 20% (v/v) acetonitrile/water. For untargeted RPLC-MS/MS analysis, half of the reconstituted samples were transferred to a new Eppendorf tube and acidified with formic acid to a final concentration of 0.1% (v/v). Meanwhile, the reconstituted supernatants were directly used for untargeted HILIC-MS/MS analysis. Samples were pooled at equal proportion to prepare global and grouped QCs for data normalization and metabolite identification, respectively.

Metabolic extracts were separated on an Accucore™ Vanquish™ C18+ UHPLC column (1.5µm; 2.1 x 150mm; ThermoFisher Scientific) for RPLC-MS/MS or a SeQuant ZIC pHILIC (5μm; 2.1 x 150mm; Supelco) column for HILIC-MS/MS. For RPLC-MS/MS, the following gradient was used: 0, 4, 8.5, 13.5, 15.5, 19, 22 min; 0.5%, 1%, 20%, 95%, 99%, 100 - 0.5% B, in which solvent A and B used was 0.1% formic acid (v/v) in water and acetonitrile, respectively. For HILIC-MS/MS, the following gradient was used: 1.5, 4, 5, 8, 11, 14, 18, 21, 25, 25.1 min; 32%, 45%, 52%, 58%, 66%, 70%, 75%, 97%, 97%, 32% B, in which solvent A and B used was 10mM ammonium acetate (pH 9.8) in water and 100% acetonitrile, respectively. Flow rates were 0.250mL/min for both chromatography columns. Compound detection was achieved by a Q-Exactive HF orbitrap mass spectrometer (ThermoFisher Scientific) operating under either positive or negative electrospray ionization mode. Samples and global QCs were analyzed by MS1 scans (m/z range: 67-1005 for positive ESI mode; 100-1005 for negative ESI mode; 120k resolution) for quantification and normalization. Grouped QC samples were analyzed by data-dependent MS2 (top 15 candidates) for metabolite identification. For metabolite annotation, database search and peak integration were performed with Compound Discoverer 3.3 (Thermo Fisher Scientific) using the public databases ChemSpider (*56*), bioCyc (*57*), HMDB (*58*), SMDPG (*59*), mzCloud/mzVault (*60*), and Cornell Proteomics Core’s in-house spectral libraries. Metabolite relative peak intensity values (data S7) were processed using normalized peak area (log_10_-transformed and Pareto-scaled) for each annotated metabolic feature for overall metabolite profile visualization using fold-change analysis, volcano plots, and PLSDA in MetaboAnalyst 6.0 (*61*).

To interrogate the metabolite functional annotations affected by microbiota conventionalization, we annotated features corresponding to well-known microbially produced or microbially-modified metabolite classes (including host-produced metabolites modified by microbes). These categories included ‘bile acids’ (primary/secondary; taurine/glycine conjugates, DCA/LCA, muricholates), ‘indole & tryptophan pathway’ (indoleacrylate/propionate/acetate/lactate, indoxyl sulfate, skatole, kynurenine/kynurenic), ‘phenylalanine/tyrosine–derived phenolics’ (p-cresol sulfate, phenylacetylglutamine, phenylacetic/propionic acids, hippurate, 4-ethylphenyl sulfate), ‘TMAO/choline axis’ (TMAO, choline, betaine, DMG), ‘SCFAs & acylcarnitines’ (acetate/propionate/butyrate proxies; C2–C5 acylcarnitines), ‘polyamines’ (putrescine, cadaverine, spermidine, spermine), ‘B-vitamins/cofactors’ (niacin/NA/NAM, riboflavin, pantothenate/CoA, biotin, thiamine, folates), ‘phenolic acids/flavonoids’ (ferulic, caffeic, vanillic, gallic, protocatechuic; catechin, quercetin), and ‘phase-II conjugates’ (sulfates/glucuronides/glycine conjugates). For each chromatography and polarity type from untargeted LC-MS/MS, we conducted paired t-tests to test whether the mean relative abundances of metabolites within each category differed significantly between AX and CV nestlings. Only categories with > 3 (the minimum required for a paired t-test) metabolites in for each assay type were tested. P-values of categories were corrected using False Discovery Rate within each assay type. For each metabolite in the ‘bile acids’ category based on results of RPLC-MS/MS in negative ion mode, we conducted Welch’s t-tests for groups with unequal variance to test for differences in peak intensities for each metabolite between AX and CV individuals. P-values from these tests were corrected for multiple comparisons using False Discovery Rate (*62*).

---

**Fig. S1.**
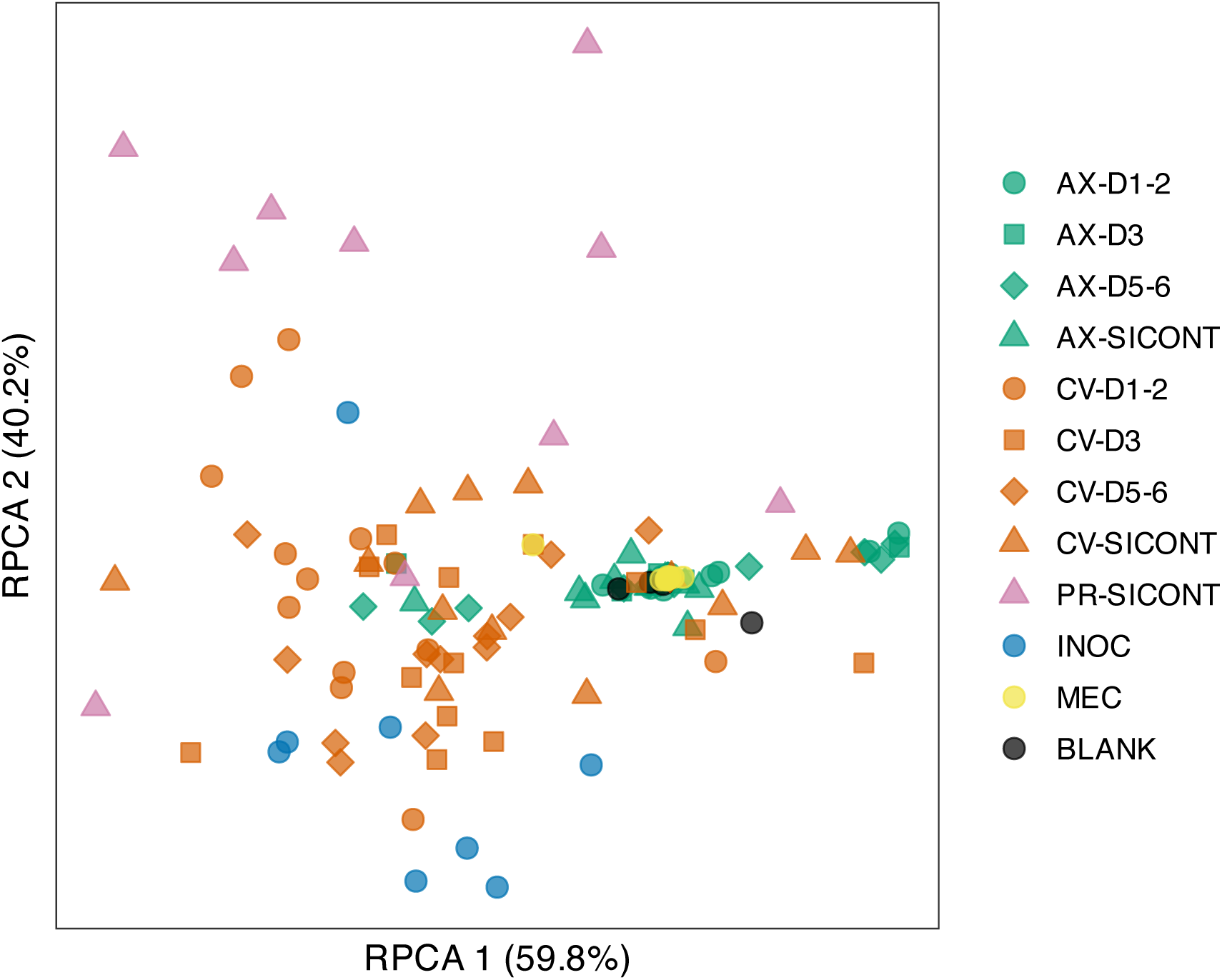
Robust Aitchison PCA (RPCA) ordination of microbiota community composition across sample types. RPCA ordination of bacterial 16S rRNA amplicon sequence variants (ASVs) shows differences in community composition across sample types, treatments, and timepoints (PERMANOVA Pseudo-F = 12.30, p = 0.001, data S2). Microbial communities of axenically raised (AX; n = 15) and conventionalized (CV; n = 13) nestlings represent bacteria from feces collected 1-2, 3, and 5-6 days post-hatch. Small intestine contents (SICONT) were collected from AX, CV, and parent-raised (PR; n = 10) nestlings at 7 days post-hatch. The meconium (MEC; n = 15) was collected from AX and CV nestlings prior to feedings or fecal inoculations (INOC; n = 8). Blank DNA extractions (BLANK; n = 16) were sequenced to control for bacterial DNA contaminants.

**Fig. S2.**
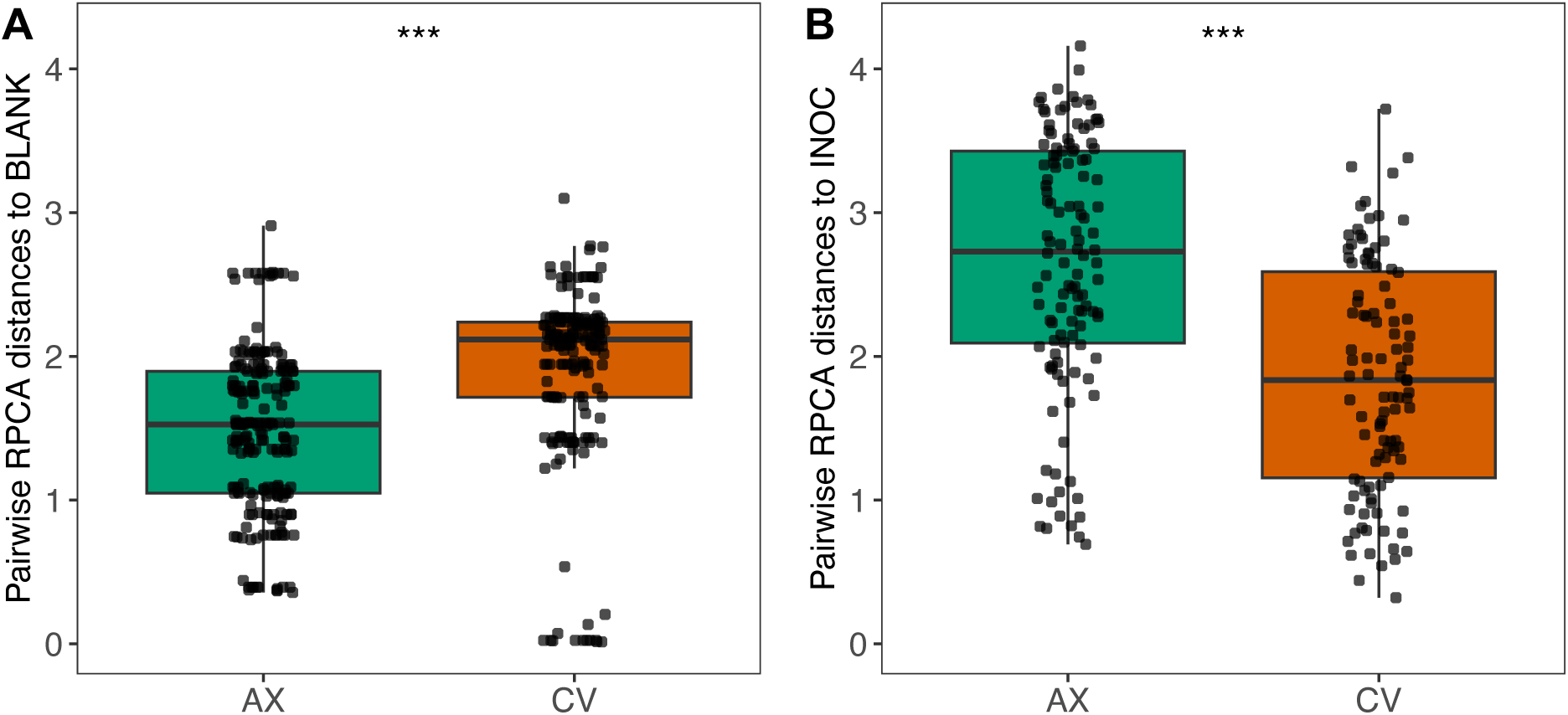
Comparison of RPCA distances across treatment groups. **(A)** Boxplot shows that axenically raised (AX; n = 15) nestlings were significantly more similar to blank extraction controls (BLANK; n = 14) compared to conventionalized (CV; n = 13) nestlings (t = 6.54; P < 0.0001). **(B)** Boxplot shows that CV nestlings were significantly more similar to microbiota of fecal inocula (INOC; n = 8) compared to AX nestlings (t = 6.40; P < 0.0001). Significance of pairwise permutation t-tests is indicated by asterisks; P ≤ 0.001 ***.

**Fig. S3.**
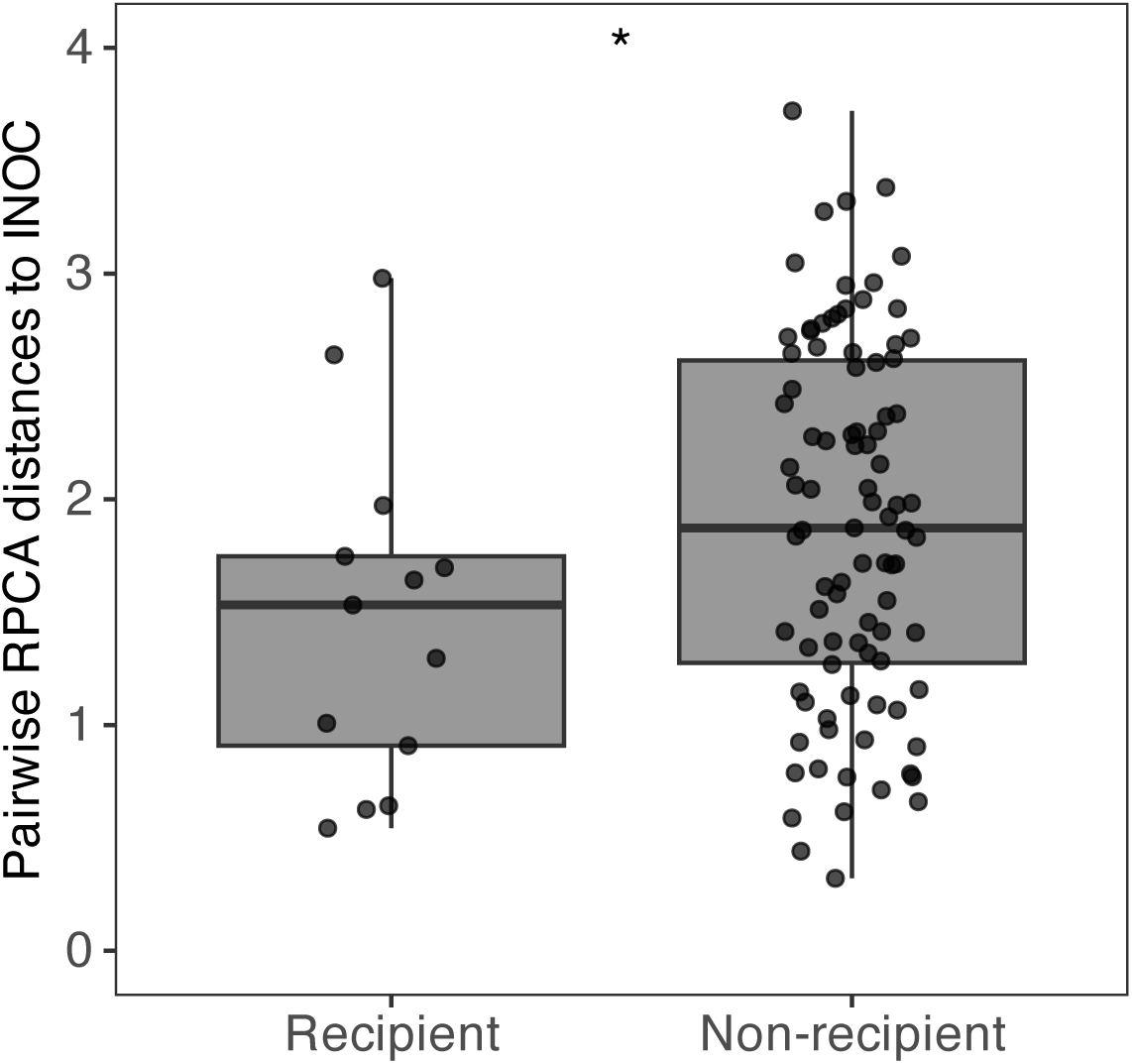
Conventionalized nestling microbiota resembles the microbial communities of the fecal inoculum received. Boxplots show that CV nestlings receiving fecal inoculations resembled their specific inocula communities significantly more than all other inocula communities (t = 2.03, P = 0.045), demonstrating efficacy of microbial transplants in recapitulating adult sparrow microbiota unique donor communities. Significance of linear mixed-effects model with individual as a random effect is indicated by asterisks; P ≤ 0.05 *.

**Fig. S4.**
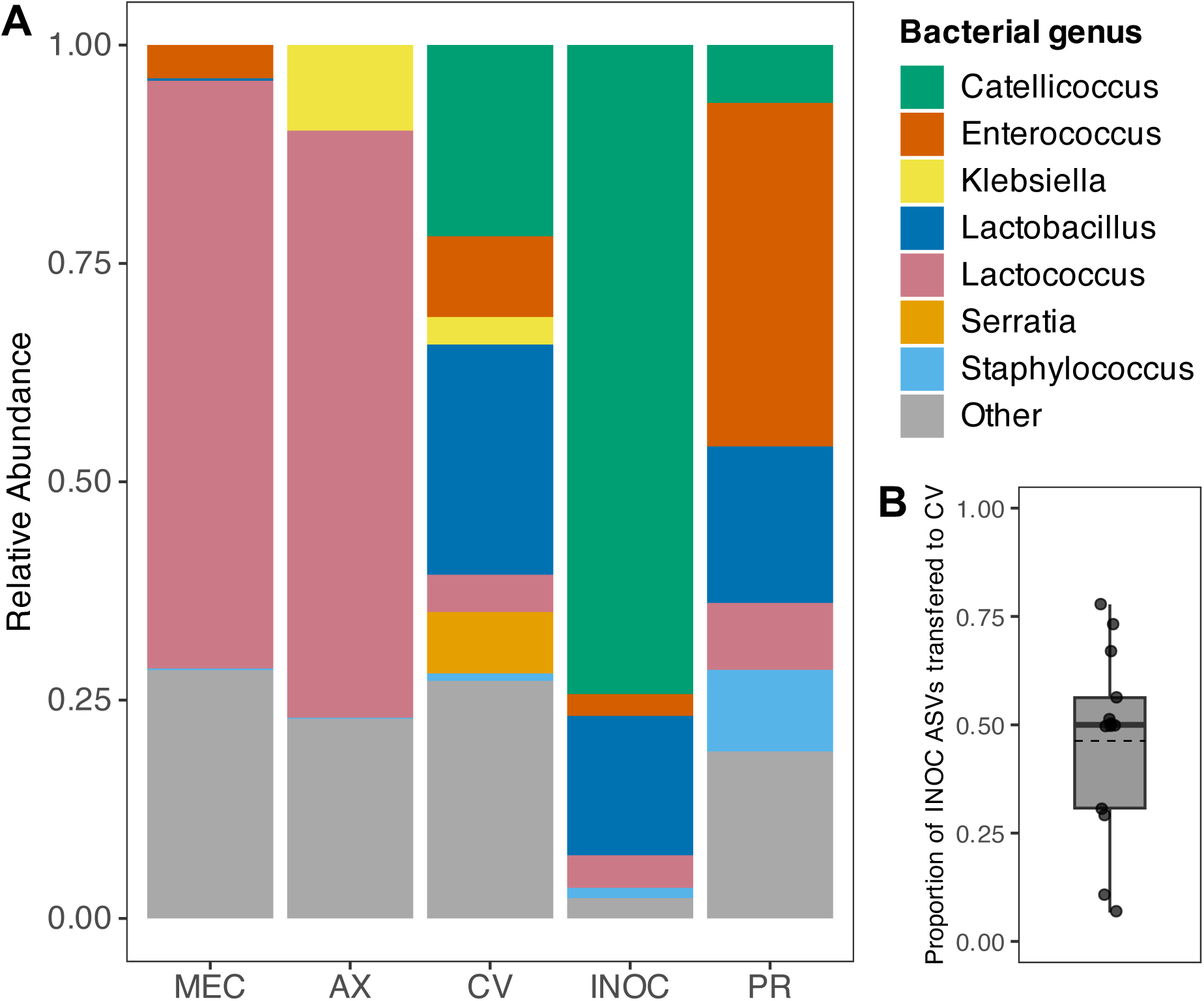
Genus-level bacterial community composition of microbiota across treatment groups. **(A)** Barplots show comparison of relative abundance of bacterial genera between meconium (MEC; n = 15) and axenically raised (AX; n = 15) nestlings 24-48 hours post hatch. Similarities in community composition between the meconium and the gut microbiota of AX nestlings demonstrate that microbial taxa in AX nestlings were present at hatch and possibly transmitted in ovo. Conventionalized (CV; n = 13) nestlings 24-48 hours post hatch were compositionally similar to inocula (INOC; n = 8) and captive parent-raised nestlings (PR; n = 10). **(B)** Boxplot shows distribution of the proportion of ASVs present in fecal inocula that were detected in the feces of their specific recipient CV nestling 24–48 hours after receiving their first inoculation, where points indicate individual CV nestlings with the dashed line indicating mean transfer efficiency.

**Fig. S5.**
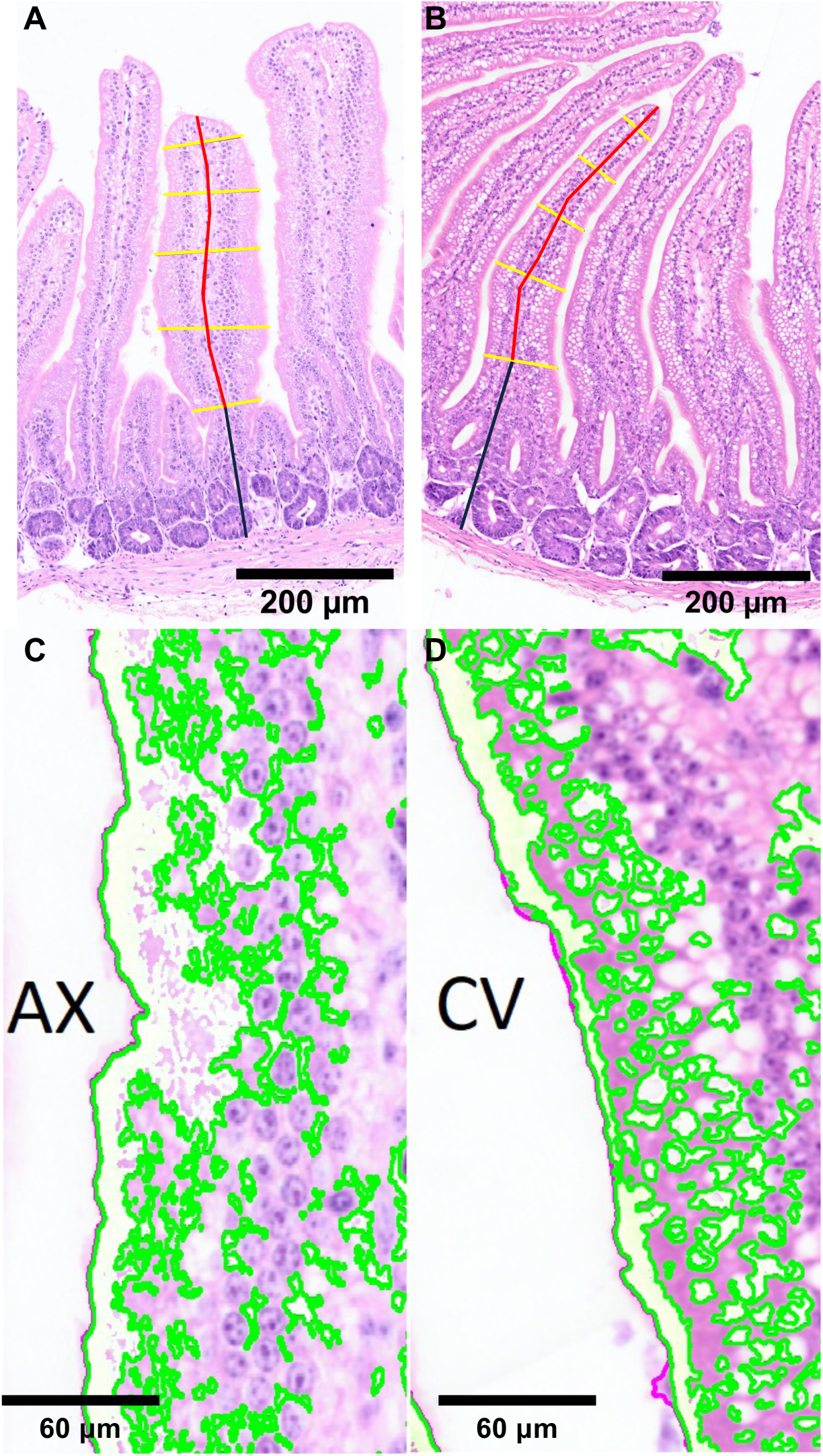
Representative images of intestinal histological quantification from experimental nestlings. Intestinal villi from **(A)** AX and **(B)** CV nestlings were quantified for length (red line), width (yellow lines), and crypt depth (black line). Intestinal goblet-cell density from **(C)** AX and **(D)** CV nestlings was quantified using HistoAnalyzer (Materials and Methods). Green overlay denotes mucosal regions containing goblet-like cells.

**Fig. S6.**
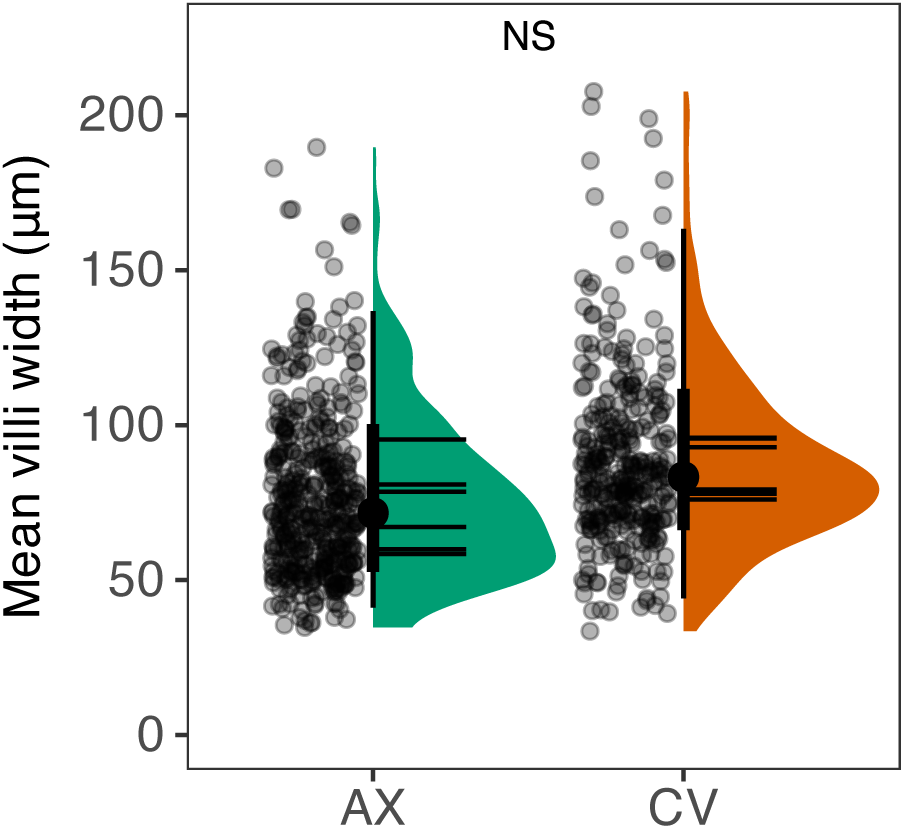
Nestling small intestine villi width was unaffected by microbiota inoculations. Violin plots of mean small intestine villi width at 7 days post-hatch show no significant difference between AX and CV nestlings (t = 1.892, P = 0.092). Points represent individual measurements, black points and bars indicate mean and standard deviation, horizontal lines within violin plots indicate individual means. NS indicates P ≥ 0.05 in linear mixed effects models accounting for sex, parentage, and individual.

**Fig S7.**
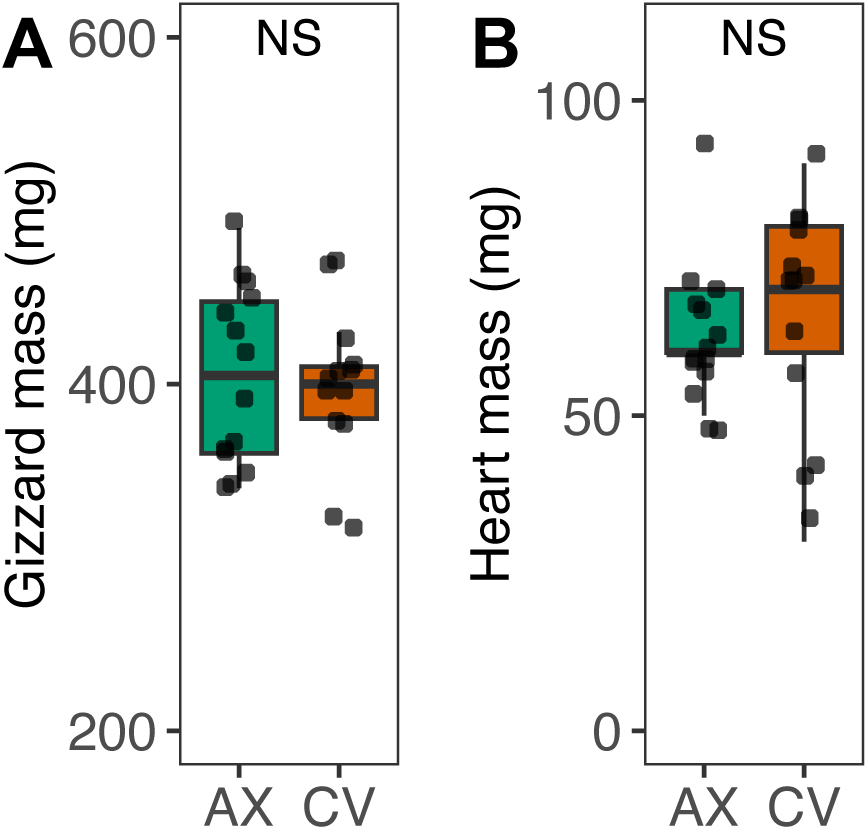
Nestling organ masses was unaffected by microbiota treatments. Boxplots show no difference between AX (n = 15) and CV (n = 13) nestlings in the mass of the **(A)** gizzard (t = 1.184, P = 0.249) or **(B)** heart (t = 0.313, P = 0.757) at days 7 post-hatch. NS indicates P ≥ 0.05 in linear mixed effects models accounting for tarsus length, sex, and parentage.

**Fig S8.**
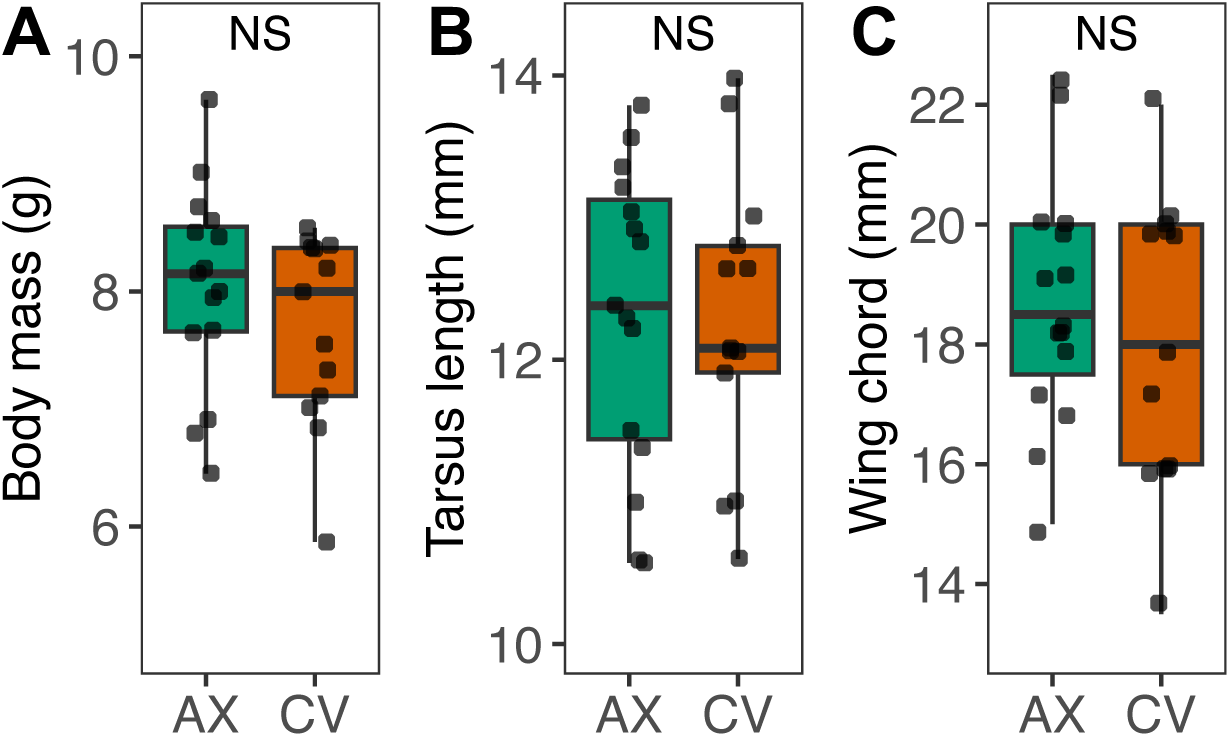
Morphometric measures of avian growth in axenically-raised and conventionalized sparrow nestlings. AX (n = 15) and CV (n = 13) nestlings exhibited no significant differences in **(A)** body mass (t = 1.99, P = 0.062), **(B)** tarsus length (t = 0.040, P = 0.969), or (C) unflattened wing chord length (t = 0.555, P = 0.584). NS indicates P ≥ 0.05 in linear mixed effects models accounting sex and parentage.

**Fig S9.**
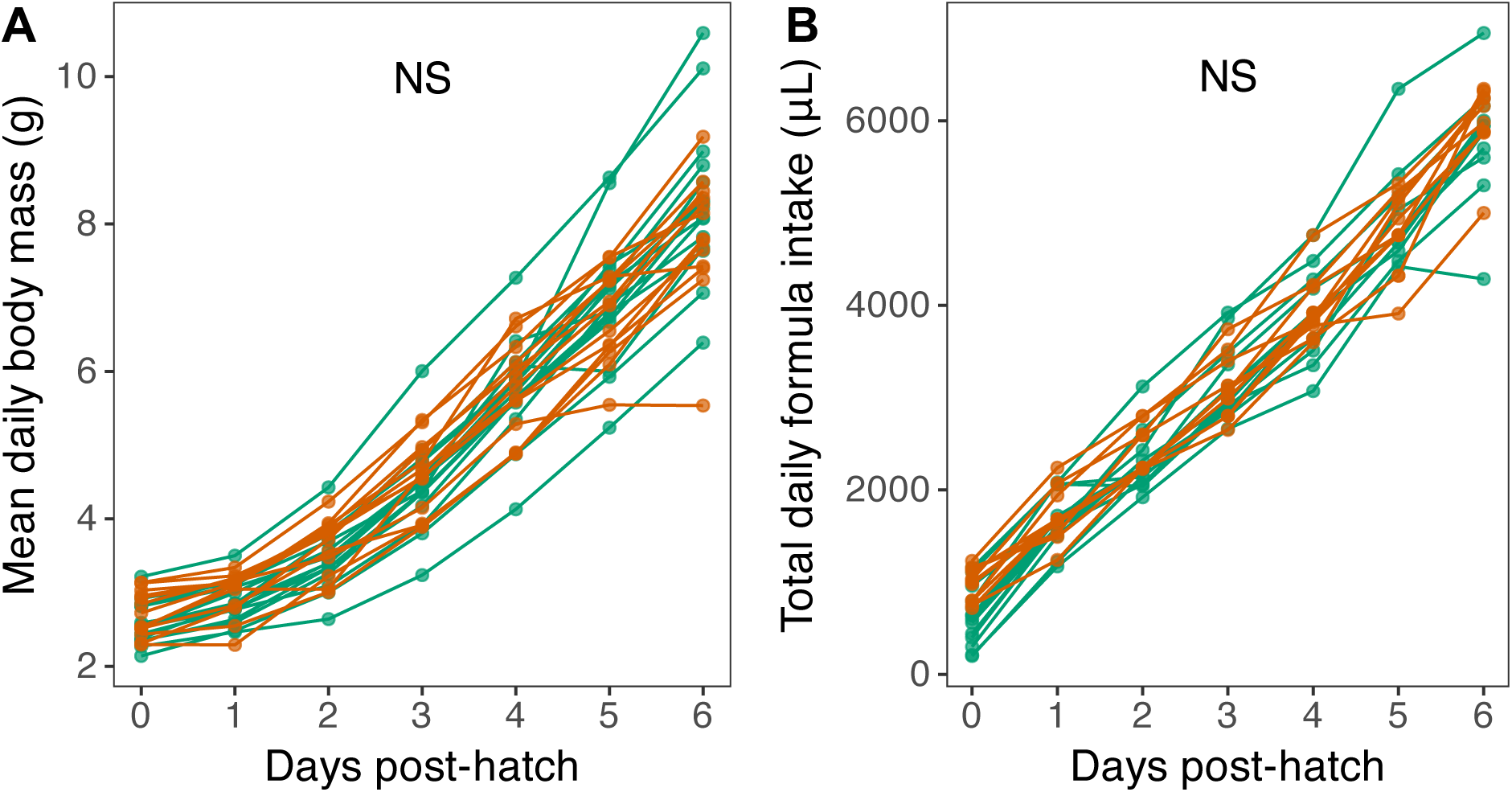
Growth rates and food intake of experimental nestlings. Line plots show AX (n = 15) and CV (n = 13) nestlings exhibited no significant differences in **(A)** mean daily body mass (t = 1.25, P = 0.218) or **(B)** total daily formula intake (t = 1.78, P = 0.082) from 0 to 6 days post-hatch. Mean daily body mass was used in A to account for variation influenced by recent feedings or measurement error. NS indicates P ≥ 0.05 in linear mixed effects models accounting sex, parentage, and individual.

**Fig S10.**
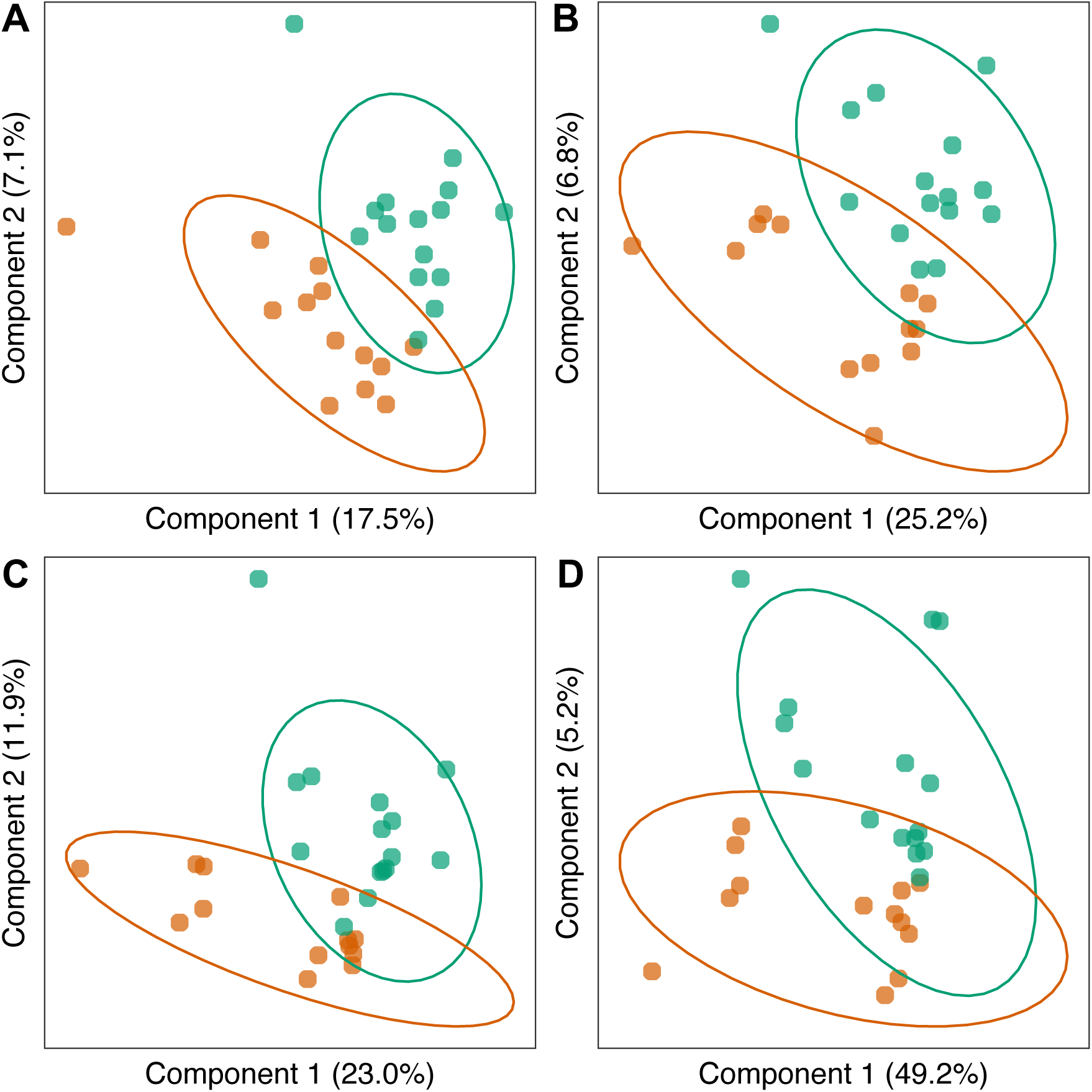
Plasma metabolite profiles differ between CV and AX birds. Visualization of PLSDA ordination showing divergent plasma metabolite profiles between AX (n = 15; green points) and CV (n = 13; red points) nestlings using **(A)** RPLC-MS/MS in negative ion mode, **(B)** RPLC-MS/MS in positive ion mode, **(C)** HILIC-MS/MS column in negative ion mode, and **(D)** HILIC-MS/MS column in positive ion mode. Ellipses represent a 95% confidence interval from group centroids.

**Fig S11.**
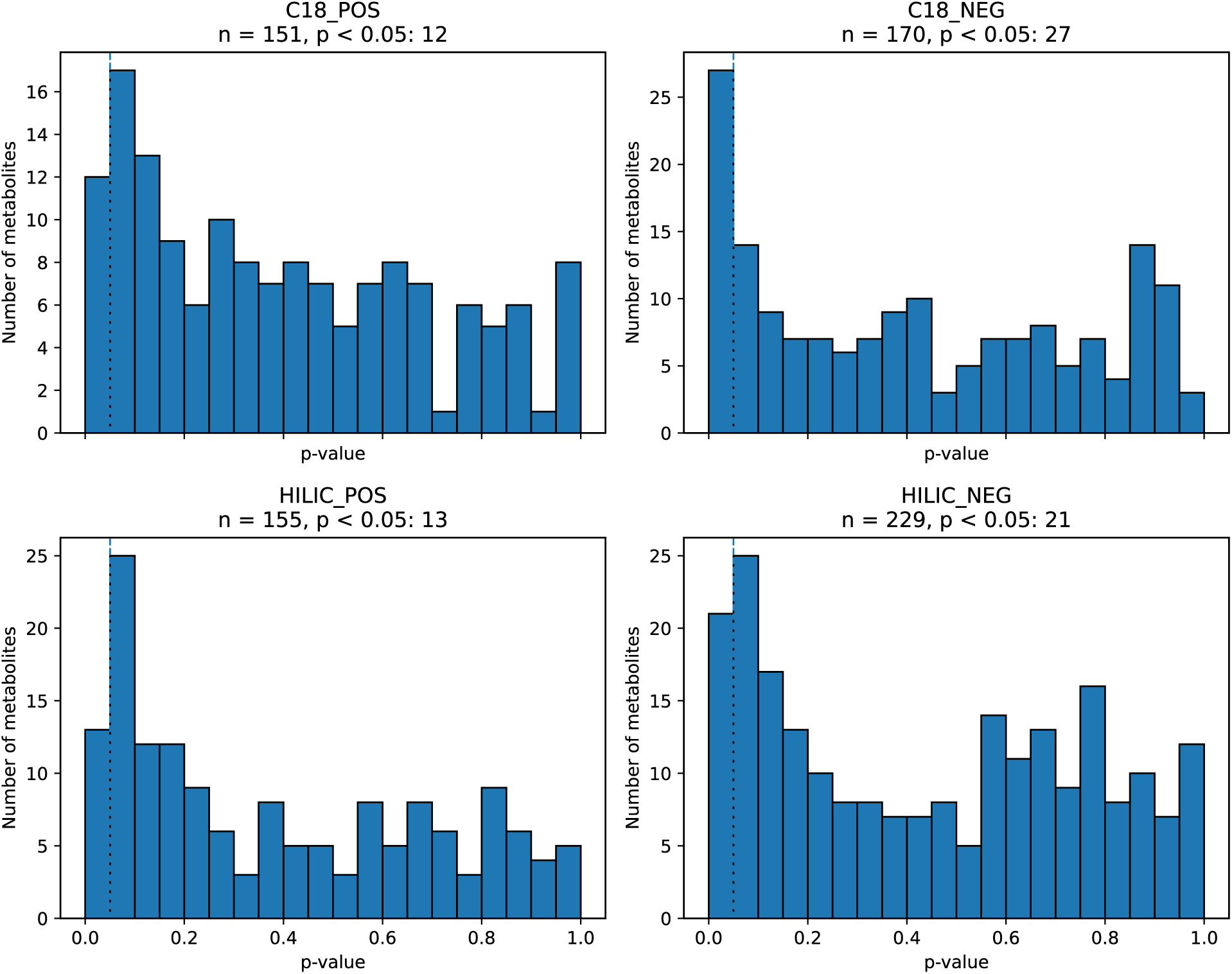
Enrichment of significant results from metabolite differential abundance testing. Histograms show distributions of p-values of linear mixed effects models used to test for differential abundances of metabolites between CV and AX nestlings using **(A)** RPLC-MS/MS in negative ion mode, **(B)** RPLC-MS/MS in positive ion mode, **(C)** HILIC-MS/MS column in negative ion mode, and **(D)** HILIC-MS/MS column in positive ion mode. Vertical dashed line denotes P = 0.05.

**Fig S12.**
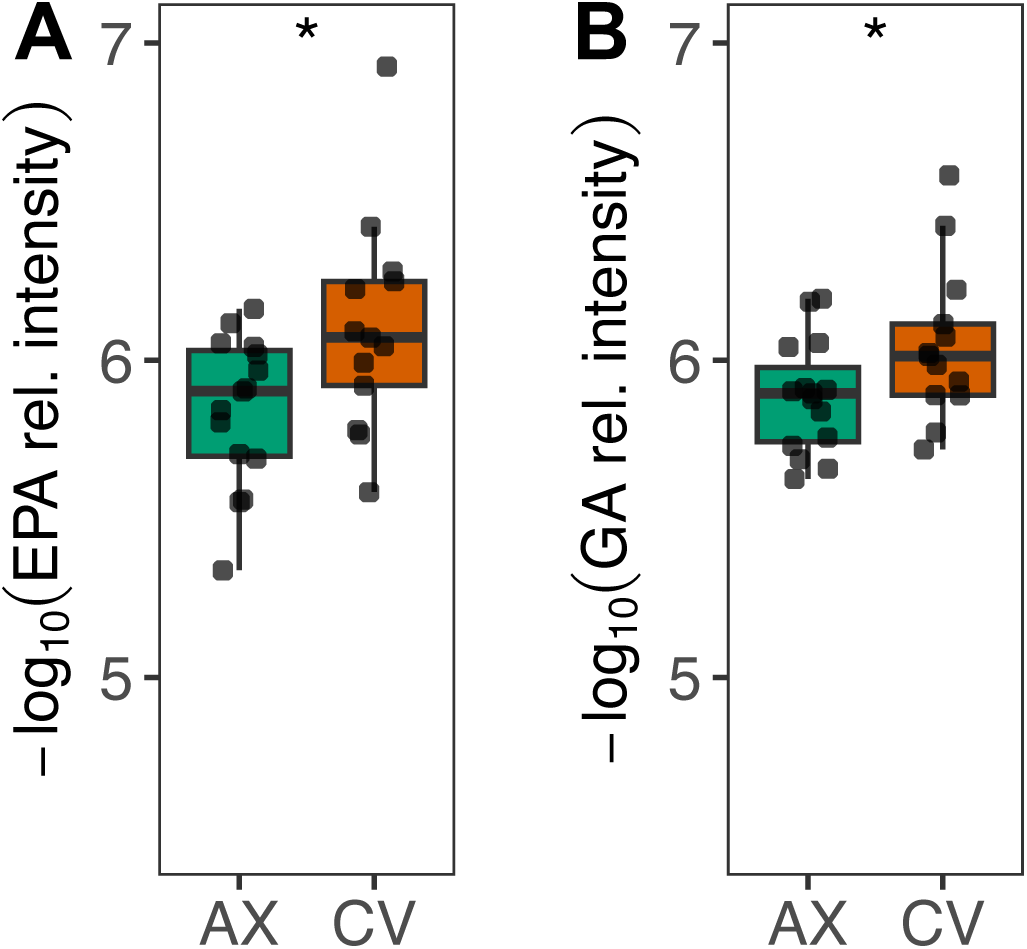
The gut microbiota affects metabolic phenotypes in songbird nestlings. Boxplots show significant elevation of log_10_-scaled relative peak intensities of metabolites associated with intestinal cellular proliferation **(A)** eicosapentaenoic acid (EPA; F = 2.942, P = 0.009) and **(B)** glyceric acid (GA; 2,3-dihydroxypropanoic acid; X2 = 2.360, P = 0.030). Significance based on linear mixed effects models accounting for parentage is indicated by asterisks; FDR-adjusted P ≤ 0.05 *.

