## Supplementary material for "Gut microbiota promote early-life digestive function in songbirds": dataS1

*Teklad Custom Diet*

# TD.200636

**Nestling Diet**

### Formula

| Formula | g/Kg |
|---|---|
| Casein | 462.3 |
| Corn Starch | 298.995 |
| Cellulose | 48.36 |
| Corn Oil | 80.0 |
| Glycine | 3.6 |
| L-Arginine HCl | 7.9 |
| L-Cystine | 3.2 |
| L-Histidine HCl, monohydrate | 1.8 |
| L-Isoleucine | 0.6 |
| L-Leucine | 1.8 |
| L-Lysine HCl | 4.1 |
| L-Methionine | 1.0 |
| L-Phenylalanine | 0.6 |
| L-Threonine | 2.2 |
| L-Tryptophan | 0.4 |
| L-Valine | 0.5 |
| Vitamin Mix, AIN-76A (40077) | 15.0 |
| Choline Chloride | 2.0 |
| Sodium Bicarbonate | 10.0 |
| Sodium Selenite (0.0445% in sucrose) | 0.45 |
| Sodium Molybdate, dihydrate | 0.09 |
| Boric Acid | 0.09 |
| Cobalt Sulfate, heptahydrate | 0.015 |
| Calcium Carbonate | 9.167 |
| Calcium Phosphate, dibasic | 26.033 |
| Sodium Phosphate, dibasic | 6.417 |
| Potassium Chloride | 6.417 |
| Magnesium Sulfate | 2.75 |
| Sodium Chloride | 3.667 |
| Ferric Citrate | 0.183 |
| Manganese Sulfate, monohydrate | 0.229 |
| Potassium Iodate | 0.009 |
| Zinc Carbonate | 0.119 |
| Cupric Sulfate | 0.009 |

### Footnote

Nesting diet patterned after the diet referenced in Physiological Zoology (1998), 71(5):561-573. Vitamins are increased to make the diet more suitable for irradiation. Silica sand (silicon dioxide) is replaced with corn starch. The last 11 items in the diet (minerals) correspond to the Fox-Briggs Mineral Mix.

### Key Features

- **Amino Acid Defined Diet**
- **Avian**
- **Nestling**
- **Vitamins Increased for Irradiation**

#### Selected Nutrient Information[1]

| | % by weight | % kcal from |
|---|---|---|
| **Protein** | 42.5 | 47.2 |
| **CHO** | 28.4 | 31.6 |
| **Fat** | 8.5 | 21.2 |
| **Kcal/g** | **3.6** | |

[1] Calculated values

[2] Protein based on N x 6.25

*Teklad Diets are designed & manufactured for research purposes only.*

### Key Planning Information

- *Products are made fresh to order*
- Store product at 4°C or lower
- Use within 6 months (applicable to most diets)
- Box labeled with product name, manufacturing date, and lot number
- Replace diet at minimum once per week

  *More frequent replacement may be advised*
- Lead time:
  - 2 weeks non-irradiated
  - 4 weeks irradiated

### Product Specific Information

- Powder
- Minimum order 3 Kg
- Irradiation available upon request

#### Options (Fees Will Apply)

- Rush order (pending availability)
- Irradiation (see Product Specific Information)
- Vacuum packaging (1 and 2 Kg)

#### Speak With A Nutritionist

- **(800) 483·5523**
- ****

#### Contact Us

*Obtain Pricing · Check Order Status*

- ****
- **(800) 483·5523**

#### International Inquiry *(Outside USA or Canada)*

- ****

#### Place Your Order (USA & Canada)

*Please Choose One*

- **www.envigo.com/teklad-orders**
- ****
- **(800) 483·5523**
- **(608) 277·2066** *facsimile*

#N/A

© 2015 Envigo

09/10/20
